# Internal limiting basement membrane inhibits functional engraftment of transplanted human retinal ganglion cells

**DOI:** 10.1101/2022.12.13.519327

**Authors:** Erika A. Aguzzi, Behnoosh Bonakdar, Marzieh Mowlavi Vardanjani, Elizabeth Kimball, Stella Mary, Kevin Y. Zhang, Jian Du, Arumugam Nagalingam, Sarah Quillen, Shreya Hariharakumar, Harry A. Quigley, Donald J. Zack, James T. Handa, Thomas V. Johnson

## Abstract

Optic neuropathies cause irreversible vision loss as retinal ganglion cells (RGCs) die. Transplantation of pluripotent stem cell (PSC)-derived RGCs offers one potential therapeutic avenue to restore vision in patients suffering from optic neuropathy if the donor neurons survive long-term in the recipient eye and develop synaptic connections within the retinal inner plexiform layer (IPL) and subcortical visual centers (*1*). Thus far, attempts at intravitreal RGC transplantation have been hampered by sequestration on the epiretinal surface without engraftment into the retinal parenchyma. In mouse retinal explant cultures, enzymatic digestion of the retinal internal limiting membrane (ILM) promotes migration of transplanted RGCs into the recipient retina (*2*). Herein, we examined donor RGC survival and engraftment in living, immunosuppressed mice, rats, and rhesus macaques and in post-mortem human retinal explant cultures. Using 3 separate human PSC lines and 3 independent methods of ILM disruption, we demonstrate that the ILM is a barrier to the retinal engraftment of intravitreally delivered human PSC-derived RGCs. ILM disruption is associated with greater donor RGC survival over 2-8 weeks and enables migration of donor neuronal somas into the endogenous RGC layer where cells elaborate dendrites into the IPL and extend axons that follow the course of the endogenous retinal nerve fiber layer into the optic nerve head. Critically, ILM disruption enables donor RGCs to synaptically integrate into IPL circuits, conferring light responsivity. These findings have important implications for enabling neuronal replacement therapies to restore vision in patients with optic neuropathy.

**SIGNIFICANCE STATEMENT:** Retinal ganglion cell (RGC) replacement and optic nerve regeneration through transplantation of stem cell-derived RGCs holds potential for restoring vision lost to optic neuropathies. Here we demonstrate that intravitreally transplanted human RGCs laminate the epiretinal surface without projecting neurites into the retinal parenchyma. However, enzymatic, developmental and surgical disruption of the internal limiting membrane not only improves graft survival, but also enables structural and functional engraftment, with dendrites that stratify the inner plexiform layer, axons that grow into the optic nerve head, and acquired responsivity to light. These observations identify a translatable approach to enable transplantation-based RGC replacement for the treatment of optic neuropathy.

## INTRODUCTION

Optic neuropathies can cause permanent vision loss as retinal ganglion cells (RGCs) die and their axons, which convey visual information from the eye to the brain via the optic nerves and tracts, degenerate. Glaucoma is the most prevalent optic neuropathy, alone affecting more than 80 million people, and ranks as the world’s leading cause of irreversible blindness (*3*). As the mammalian retina lacks inherent regenerative capacity (*4–6*), restoring lost vision in patients with optic neuropathy will require regenerative approaches that replace functional RGCs within the anterior visual pathway (*1*).

Neuronal transplantation is a promising approach to restore function in neurodegenerative disorders (*1, 7–9*). Subretinal transplantation of photoreceptors improves vision in animal models of retinal degeneration (*10–12*) and is nearing human clinical trial. However, transplanting RGCs has been more challenging, owing to their more complex neurophysiology, reception of afferent input from bipolar and amacrine cells within complex and diverse inner plexiform layer (IPL) circuits, and long-distance axonal projections to the brain. Key advancements over the past 15 years have described several efficient methods for generating RGCs from human pluripotent stem cells (PSCs) (*13–15*) and have identified numerous molecular pathways that can be manipulated to enable the long-distance regeneration of endogenous RGC axons from the retrobulbar optic nerve, past the chiasm, and into key subcortical visual targets (*16–29*). In some cases, modulation of these pathways has even improved rudimentary visually guided behaviors in mice (*16, 30*). These discoveries substantiate the feasibility of postsynaptic brain connectivity by repopulated RGCs is feasible.

Nonetheless, rigorous demonstrations of synaptogenesis by transplanted RGCs are lacking, and methods to augment this process require further development. Previous reports have suggested rare instances of functional integration by transplanted primary neonatal mouse RGCs (*31*), murine PSC-derived RGCs (*32, 33*), and human PSC-derived RGCs (hRGCs) (*34*) in recipient retinas. However, prior studies have not definitively ruled out intercellular material transfer as a potential confound that produces erroneous labeling of endogenous RGCs with donor reporter molecules and subsequent misclassification of RGC origin (*35, 36*). Indeed, there is a precedent for such misidentification, as intercellular material transfer was found to contribute to flawed conclusions about the engraftment efficiency of photoreceptors transplanted into the subretinal space of rodents (*35–39*). Moreover, among the relatively few RGCs that survive transplantation (typically <1%) (*7, 9, 21, 31, 40*), only a small fraction of RGCs purportedly engraft into the retina by migrating into the host retinal ganglion cell layer (RGCL) and stratifying dendrites into the IPL, where synaptogenesis with host bipolar and amacrine cells could occur.

Donor neurons encounter significant barriers to retinal incorporation. Intravitreal injection of a cell suspension disperses the graft into an acellular cavity devoid of direct metabolic support. The internal limiting membrane (ILM), the basement membrane at the vitreoretinal interface, represents a key obstacle that limits retinal transit of drugs, gene therapy vectors, and cells into the neural retina (*2, 41–44*). The ILM is composed of extracellular matrix (ECM) proteins including collagen (type IV, XVIII), laminin, fibronectin, and nidogens 1 and 2 that are deposited atop a layer of Müller glial endplates (*46*). During development, integrin-mediated ILM recognition by RGCs is required for appropriate RGC polarization, migration, and dendritic lamination (*45–48*), and ILM-astrocyte interactions modulate glial migration that guides subsequent retinal vascular development (*49, 50*). However, in adulthood the ILM is not only dispensable, but can serve as a scaffold for the pathological proliferation of myofibroblasts, retinal pigment epithelium, and glial cells. Indeed, surgical ILM removal is used to relieve vitreomacular traction and can improve vision in patients with macular holes and epiretinal membranes, supporting the feasibility of disrupting or removing the ILM in human patients to augment cell therapy (*51*). It has long been speculated that the ILM might impede retinal integration of intravitreally transplanted RGCs (*52, 53*). Here, we evaluate enzymatic, developmental, and surgical approaches of ILM disruption for promoting the structural and functional engraftment of transplanted hRGCs in immunosuppressed mice, rats, and non-human primates *in vivo* and in human postmortem organotypic retinal explant cultures.

## RESULTS

### ILM disruption through proteolytic and developmental approaches in rodents

We have demonstrated that proteolytic ILM digestion with pronase-E increases donor hRGC retinal neurite growth in murine organotypic retinal explants by ≥40-fold without causing detectable neural or glial toxicity (*2*). To determine whether this approach is efficacious *in vivo*, we intravitreally administered pronase-E or balanced salt solution (BSS) as vehicle in adult wild type C57BL/6J mice and Wistar rats (Fig 1A, F). In mice, we found that the concentration of pronase-E that optimally digested the ILM was partially lot dependent (1.25 milliunits for lot SLCK2824 and 0.075 milliunits for lot SLCL6675, see *Methods*). Intravitreal pronase-E concentrations ≤1.0 milliunits caused no histological abnormalities in the retina, whereas concentrations ≥2 milliunits were associated with visible preretinal and intraretinal hemorrhage (Fig 1B), presumably due to digestion of retinal vasculature basement membranes. We found similar effects following intravitreal collagenase administration *in vivo* in rats (*54*). Concentrations ≥6 milliunits caused severe intraocular hemorrhage and gross degradation of the neuroretina (Fig 1B). We performed similar pronase-E dose titration experiments in rats and found that the optimal dose was 2 milliunits (lot SLCL6675, Fig 1G).

**Figure 1.**
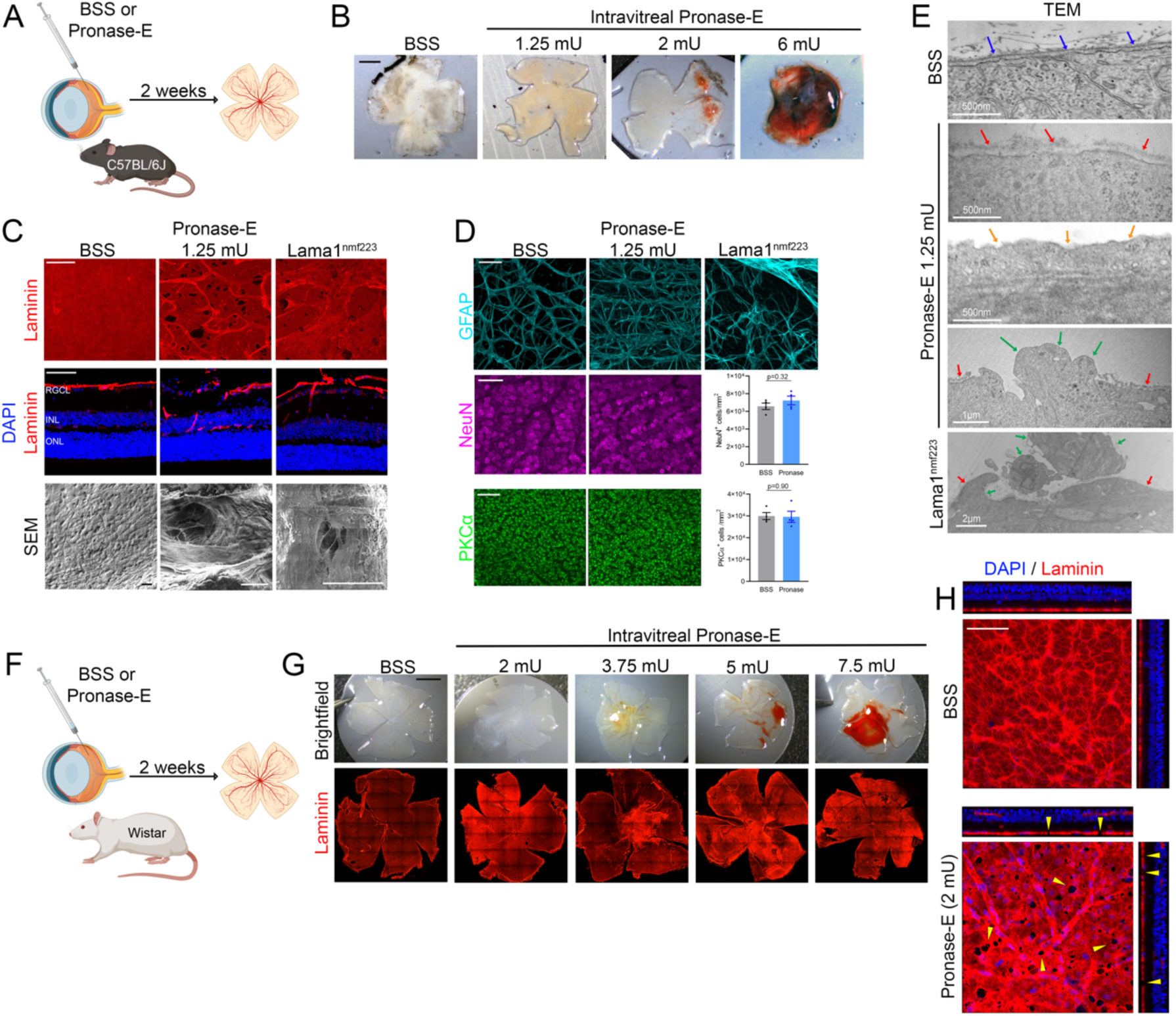
Internal limiting membrane (ILM) disruption in rodents. Pronase-E or BSS was injected intravitreally in mice (A) or rats (F) and retinae examined 2 weeks later. Retinal flat-mounts demonstrated preretinal and intraretinal hemorrhage with supratherapeutic doses of Pronase-E (B, mice; G, rats; scalebars, 1mm). (C) Immunofluorescence (IF) and scanning electron microscopy (SEM) demonstrated ILM defects in Lama1^nmf223^ and Pronase-E-treated retinas. Top and bottom rows: *en face*; middle row: cryosection. Scalebars, IF = 50µm, SEM = 20µM. (D) IF and quantification of retinal glia and inner retinal neurons. Scalebars, 50µm. (E) Transmission electron microscopy of the inner retina. Arrows: blue, intact ILM; red, disrupted ILM; orange, absent ILM; green, glial protrusion. H) Maximal projections and orthogonal projects of z-stack in rat retinae. Arrowheads, ILM breaks. Scalebars, 50µm. mU, milliunits.

We evaluated the effects of intravitreal pronase-E on the ILM and underlying retina in several ways. Laminin in BSS-treated retinas exhibited uniform immunoreactivity across the ILM surface, whereas the ILM of pronase E-treated retinas contained diffuse full-thickness round defects ranging in size up to 400 µm^2^ (Fig 1C, H, Supp Fig S1A,B). Ultrastructurally, the ILM appeared dense, continuous, and uniform in BSS-treated eyes, with Müller glial endplates abutting the basement membrane (Fig 1E). In contrast, pronase-E was associated with ILM thinning, irregularity, and frank discontinuities through which glial cells occasionally protruded (Fig 1E). The ILM surface was continuous and uniform in BSS-treated eyes, whereas pronase-E induced sizable craters and holes in the ILM when imaged with scanning electron microscopy (SEM, Fig 1C).

To assess the effects of pronase-E on resident inner retinal cells, we evaluated neuronal and glial morphology using immunofluorescent confocal microscopy. GFAP immunofluorescence in the retinal nerve fiber layer (RNFL) demonstrated modest hypertrophy of astrocytes and Müller glial endplates following pronase-E treatment in comparison to BSS (Fig 1D; Supp Fig S1A,B), but did not induce glial cell death as we observed previously with other proteolytic enzymes or high doses of pronase-E (*2*). Neither RGCL neuron density (RGCs and displaced amacrine cells labeled with NeuN) nor rod bipolar cell density (labeled with PKCα) were affected by pronase-E administration (Fig 1H,I). Together these data show that intravitreal pronase-E administration can be effectively titrated to digest and produce full-thickness defects in the ILM without causing inner retinal cell death, although with modest reactive gliosis.

We considered that proteolytic ILM digestion may confer occult off-target effects on the retina not identifiable on routine histology, which could affect RGC engraftment. Therefore, we evaluated an independent, complementary model of ILM disruption. *Lama1^nmf223^* mice (C57BL/6J background) harbor a missense mutation in *Lama-α1,* which reduces laminin-α1 protein interactions at the N-terminal domain where receptor binding and polymerization motifs are located. These mice exhibit developmental ILM defects associated with ectopic migration of retinal astrocytes into the vitreous cavity, followed by the development of patchy epiretinal gliovascular membranes, as previously characterized (*49, 50*). We reasoned that ILM defects, which permit astrocyte extravasation from the retina into the vitreous cavity, may also enable migration of intravitreal donor RGCs into the retina.

Before transplanting RGCs, we further characterized the ILM disruption exhibited by these mice. Like pronase-E-treated retinas, the ILM of *Lama1^nmf223^* mice had full-thickness defects (Fig 1C, Supp Fig S1C), which were generally sparser than following pronase-E administration. We also identified a preretinal gliofibrotic membrane in these mice, as has been described previously (Fig 1C, Supp Fig S1C) (*49, 50*). Local breaks in the ILM over diffuse areas of the retinal surface exposing bare Müller glial endplates, and in some instances, glial projections into the vitreous cavity were visualized ultrastructurally (Fig 1E). Pits and full-thickness holes, similar to that of pronase-treated retinas, were visualized with SEM (Fig 1E).

### Donor hRGC engraftment following ILM disruption in rodents

Prior to transplanting hRGCs *in vivo*, we compared engraftment of hRGCs within organotypic retinal explant cultures derived from adult C57BL/6J mice versus *Lama1^nmf223^* mice (Supp Fig S2A). hRGCs were differentiated and purified from human H7 hESCs as previously described and extensively characterized (*13, 40, 55*). Like prior observations of pronase-E-treated retinal explants (*2*), we observed that tdTomato (tdT) expressing hRGCs survived for ≥7 days in co-culture and that hRGCs on retinas with intact ILM tended to form discrete clusters (Supp Fig S2B). Further, hRGCs extended robust neurites laterally on the epiretinal surface, but most remained superficial to the retinal parenchyma, and specifically, to the astrocytes within the RNFL (Supp Fig S2C). A subset of hRGCs exhibited structural engraftment, with somas localized to the RGCL and neurites that grew deeper into the host neural retina (Supp Fig S1D). Transplantation onto *Lama1^nmf223^* retinas was associated with greater donor hRGC survival (Supp Fig S2E) and neurite extension into the recipient tissue (Supp Fig S2F).

We next evaluated the effect of ILM disruption on donor hRGC engraftment *in vivo* in mice. We tested both ILM disruption models in parallel by transplanting 4×10^5^ tdT^+^ hRGCs derived from H7 hESCs into immunosuppressed mice and assessed retinal integration 2 weeks later (Fig 2A,B). hRGCs survived in the majority of eyes (75% of BSS-treated eyes (n=12/16); 80% of pronase-treated eyes (n=32/40); and 95% of *Lama1^nmf223^* eyes (n=45/50)). As in retinal explants, hRGCs transplanted *in vivo* tended to clump in the presence of an intact ILM, and survival and cell dispersion were higher when the ILM was disrupted (Fig 2C, D). Remarkably, RGCs frequently extended a lengthy axon that was directly linearly towards the optic nerve head (ONH, Fig 2C). The absolute number of surviving hRGCs remained generally <1% of the number injected (Fig 2D), commensurate with prior reports (*7, 9, 31, 32, 34*).

**Figure 2.**
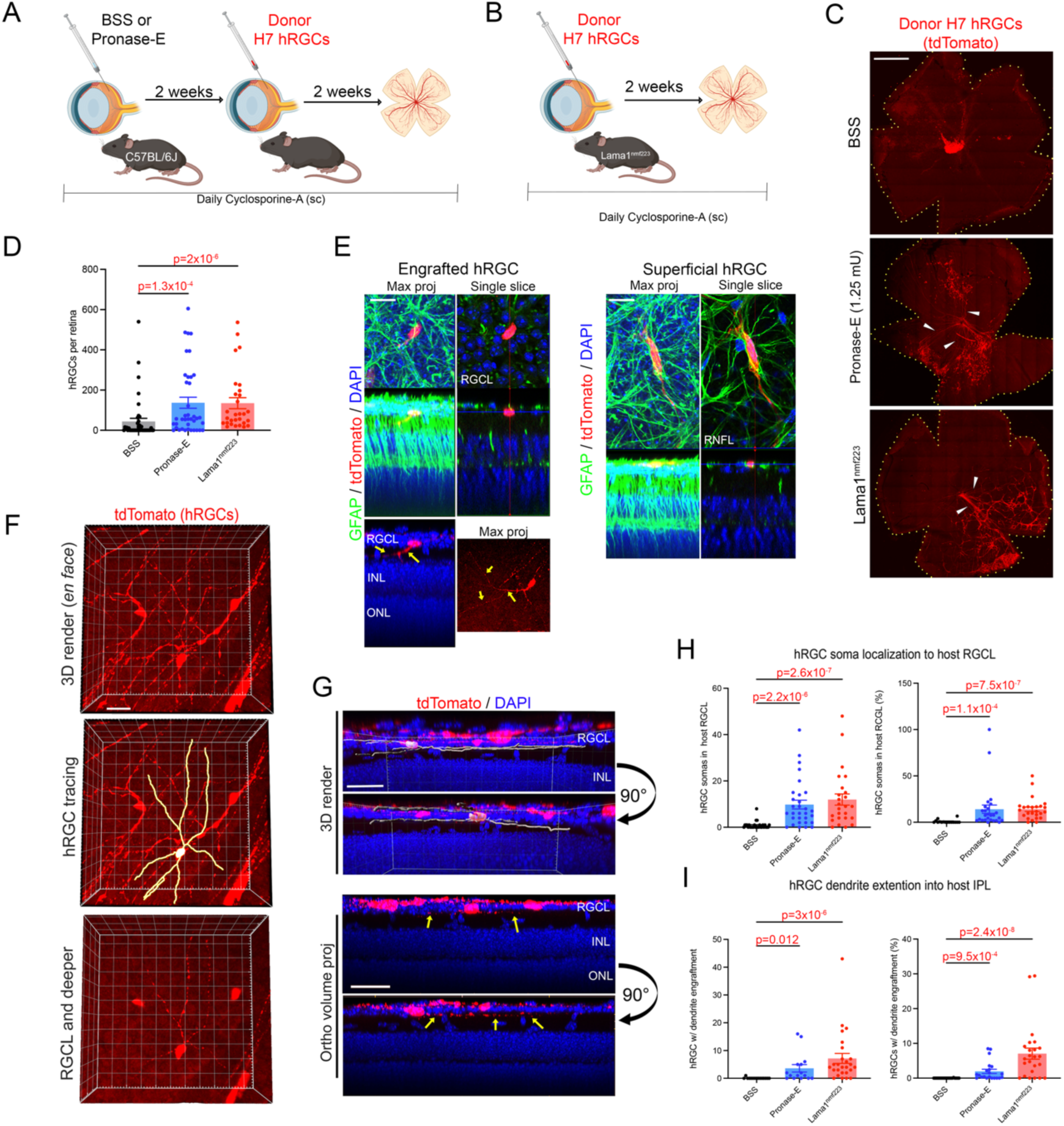
Structural H7-hRGC engraftment in mice. H7 hRGCs were intravitreally injected in immunosuppressed wildtype mice following pronase-E treatment (A) or directly into Lama1^nmf223^ mice (B). (C) Retinal flat-mounts demonstrating hRGC survival and localization 2 weeks after transplantation. Arrowheads, hRGC axon extension towards optic nerve head. Scalebar, 1mm. (D) Quantification of hRGC survival per eye. (E) Confocal images demonstrating structural engraftment or engraftment failure (superficial hRGC). Arrows, hRGC dendrites within the inner plexiform layer (IPL). Scalebar, 20µm. (F) 3D rendering of confocal z-stack showing an engrafted hRGC, traced in the middle panel. Scalebar, 30µm. (G) Alternate view of the hRGCs shown in panel F (hRGC tracing in grey). Arrows, hRGC dendrites in the IPL. Scalebar, 20µm. Quantification of absolute and relative hRGC soma (H) and dendrite (I) engraftment.

We next characterized the structural engraftment of hRGCs using two primary outcomes: localization of hRGC somas to the endogenous RGCL and lamination of hRGC dendrites within the endogenous IPL. For these studies, we carefully assessed 3D renderings of high-resolution confocal z-stacks of flat-mounted retinas, since transverse histological sections would capture only fragments of hRGC dendrites oriented in the orthogonal plane. Within the host retina, most hRGCs were localized to the epiretinal surface, wherein the hRGC soma was superficial to the RGCL and astrocytic GFAP processes within the RNFL (Fig 2E). In contrast, some hRGCs migrated into the RGCL, with a soma that was coplanar with the endogenous RGCL and deep to the astrocytes of the RNFL (Fig 2E). Negligible somal migration into the RGCL occurred in BSS-treated eyes, whereas the absolute number and percentages of surviving hRGCs that engrafted were significantly higher in pronase-treated or *Lama1^nmf223^* eyes (Fig 2H).

Individual hRGCs with neurites within the retinal parenchyma were traced and the localization of the neurites was assessed in a retinal layer-specific manner. Unlike our prior observations after transplanting hRGCs onto organotypic retinal explants, wherein donor hRGC neurites appeared to grow throughout the retina in a nonspecific manner (*2*), hRGC dendrites transplanted *in vivo* strikingly targeted the IPL for lamination (Fig 2F, G, Supp Videos S1-2; https://youtu.be/KCRGL8-1X20 and https://youtu.be/eBzE08K3k7E). In BSS treated retinas, we observed rare hRGCs which had extended dendrites into the IPL, whereas this phenomenon occurred readily in pronase-E-treated and *Lama1^nmf223^*eyes (Fig 2I).

These experiments were replicated by transplanting hRGCs from 2 additional independent human PSC lines (H9 hESCs and EP1 hiPSCs (*56*), derived as above for H7 hESCs (*13*)) into C57BL6/J mice with BSS or pronase-E treatment or Lama1^nmf223^ mice (Supp Fig S3,4). Results were generally similar: more H9 hRGCs survived with ILM disruption, and in that context both hRGC derivations achieved striking increases in somal and dendritic engraftment. These results substantiate that the barrier effect of the ILM is not specific to RGCs derived from a single source.

To determine whether the effects of ILM disruption on donor hRGC engraftment may be species-specific, we transplanted H7 hRGCs in rats following intravitreal injection of either BSS or pronase-E (Fig 3A). As in mice, there was greater survival, somal engraftment, and dendrite lamination within the IPL in eyes treated with pronase-E as compared to BSS-treated eyes with intact ILM (Fig 3B-D).

**Figure 3.**
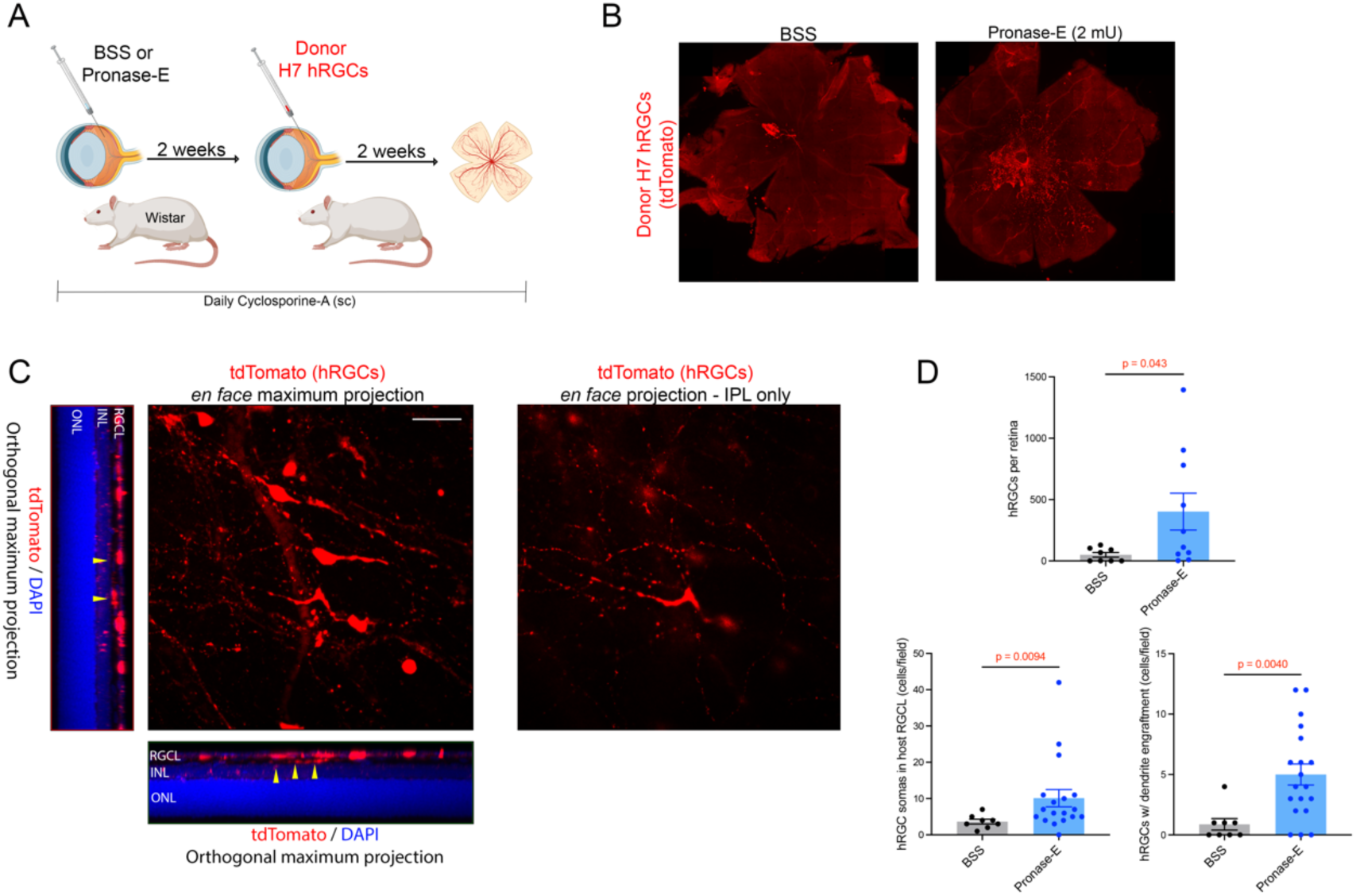
Structural hRGC engraftment in rats. H7 hRGCs were intravitreally injected in immunosuppressed Wistar rats following pronase-E treatment (A). (B) Retinal flat-mounts demonstrating hRGC survival and localization 2 weeks after transplantation. Scalebar, 1mm. (C), *En face* and orthogonal projections of confocal z-stack showing an engrafted hRGCs (left) and *en face* projection through the inner plexiform layer only, demonstrating hRGC dendrite projections. Scalebar, 20µm. (D) Quantification of hRGC engraftment.

While the level of hRGC survival observed was comparable to past reports, increasing graft survival rates will be critical to the future translatability of neuronal transplantation. We wondered whether cyclosporine-A immunosuppression in wildtype mice may have been insufficient for preventing xenograft rejection and contributed to suboptimal hRGC survival. Thus, we compared transplantation of hRGCs in cyclosporine-immunosuppressed C57BL/6J mice to NOD-SCID-gamma (NSG) mice, which carry mutations in *Prkdc* and *Il2rg* leading to the absence of mature T cells, B cells, and NK cells (*57*). There was no improvement in hRGC survival in mice lacking an adaptive immune system (Supp Fig S5), suggesting that breakthrough adaptive immune rejection is negligible in mice systemically immunosuppressed with cyclosporine-A. hRGC death at this relatively early time point is therefore likely to be related to other factors, such as innate immune activity (i.e. by microglia or macroglia) and/or paucity of supportive factors in the microenvironment.

### Donor hRGC engraftment in an ocular hypertensive glaucoma mouse model

Given our intent of translating RGC repopulation to optic neuropathy, we next transplanted hRGCs in a mouse model of ocular hypertensive glaucoma (Fig 4A). Lama1^nmf223^ mice underwent intracameral microbead injection into one randomly selected eye and BSS (vehicle) contralaterally. Intraocular pressure (IOP) rose 24h following microbead injection by 10-20mmHg, consistent with our prior work (Fig 4B). hRGC transplantation was performed 8 weeks after experimental glaucoma induction, when IOP had returned to normal. Ocular hypertensive glaucoma was associated with a >15-fold increase in donor hRGC survival as compared to naïve, healthy eyes (Fig 4C,D). In addition, engraftment of hRGCs into the retina was robust in Lama1^nmf223^ mice and, interestingly, the glaucomatous microenvironment was associated with significant increases in hRGC somal localization to the endogenous RGC layer and dendrite stratification within the IPL (Fig 4D-F). These data show that hRGC transplantation is not only feasible in glaucomatous retinas, but also yields greater survival and engraftment than in healthy retinas.

**Figure 4.**
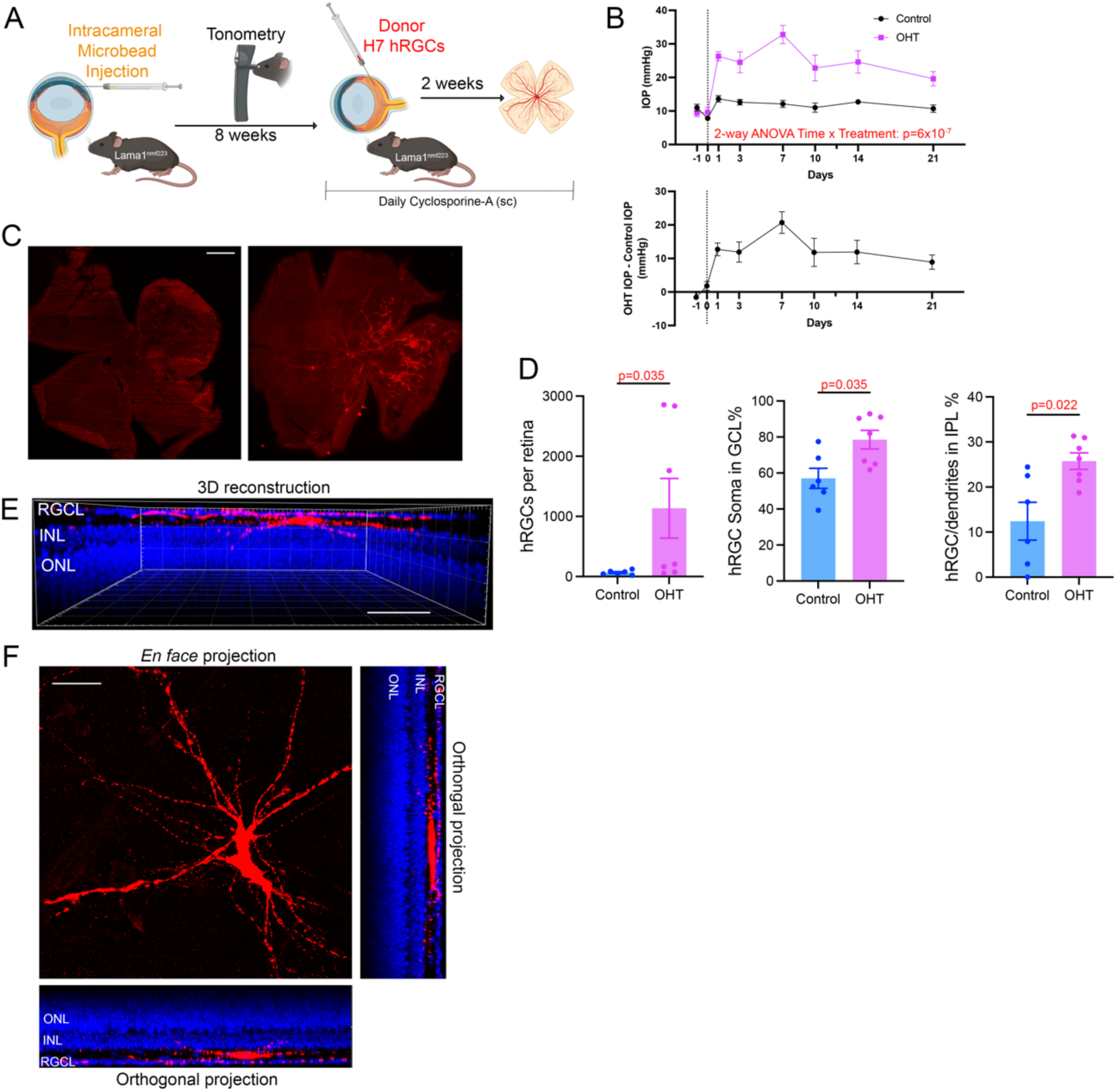
Structural hRGC engraftment in a mouse model of ocular hypertensive glaucoma. (A) Immunosuppressed Lama1^nmf223^ mice underwent intracameral microbead injection to elevate intraocular pressure (IOP) followed by H7 hRGC transplantation 8 weeks later. (B) IOP elevation following intracameral microbead injection. (C) Retinal flat-mounts demonstrating hRGC survival and localization 2 weeks after transplantation. Scalebar, 1mm. (D) Quantification of hRGC survival and engraftment following transplantation. (E) 3D rendering of confocal z-stack showing an engrafted hRGC. Scalebar, 20µm. (F) *En face* and orthogonal projections of confocal z-stack showing the engrafted hRGC from panel E.

### Transplantation of hRGCs onto human organotypic retinal explants

The translational potential of RGC transplantation necessitates ultimately achieving engraftment into human retinas. The structure and composition of the ILM varies among species, and as a function of age (*58*). Thus, we wondered whether the ILM also obstructs hRGCs from extending dendrites into human retinal tissue, and whether this blockade can be ameliorated with enzymatic ILM digestion. We obtained ocular tissue from human postmortem donors (ages 72y and 87y) within 24h of death, isolated their retinas, and cultured them in organotypic fashion. As above, we treated the retinal tissue with pronase-E (or BSS, vehicle) onto the inner retinal surface, and then transplanted 1.5×10^5^ hRGCs 24h later (Fig 5A). At 7d after co-culture, hRGCs survived on the surface of retinal explants and extended long neurites (Fig 5B), similar to their behavior on mouse retinal tissue. hRGC survival rates were similar in retinas pre-treated with pronase or BSS (Fig 5D). hRGCs elaborated dendrites within the IPL at a significantly higher rate when the ILM was permeabilized as compared to intact (Fig 5C,E). These data demonstrate that the ILM inhibits hRGC engraftment into human neural retinas and supports the translational premise of circumventing the ILM for hRGC transplantation.

**Figure 5.**
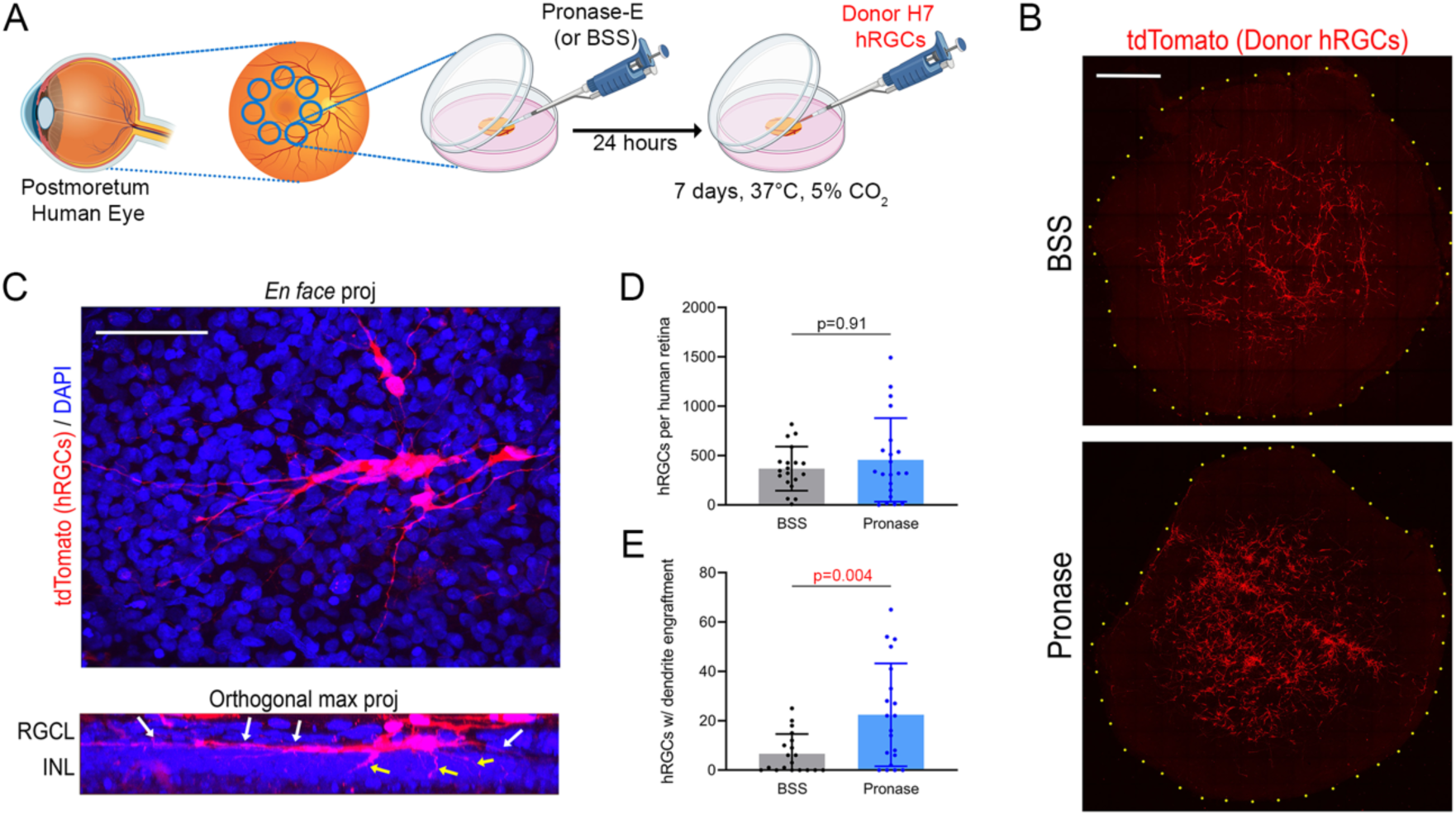
Structural hRGC engraftment in postmortem human organotypic retinal explants. (A) Retinal explants were generated from the maculae of human donor eyes (4 eyes of 2 donors), treated with BSS or pronase-E, and cultured for 1 week. hRGCs were applied on 24h after dissection. Dots outline the edge of the explant. (B) Retinal flat-mounts demonstrating hRGC survival and localization after 1 week of co-culture. Scalebar, 1mm. (C) *En face* and orthogonal projections of confocal z-stack showing an engrafted hRGC. Arrows, dendrites within the inner plexiform layer and inner nuclear layer. Scalebar, 20µm. Quantification of hRGC survival (D) and dendrite engraftment (E).

### Transplantation of hRGCs into non-human primates

To develop a translatable model for hRGC transplantation, we performed experiments using immunosuppressed rhesus macaques (non-human primates, 3-4y old, N=4, Fig 6A). Two animals underwent unilateral induction of experimental ocular hypertensive glaucoma by gonioscopic laser photocoagulation of the trabecular meshwork and 2 remained naïve with healthy eyes. IOPs of glaucoma eyes were significantly elevated by 3 weeks after the initial laser treatment and remained high (30-60 mmHg) for the duration of the experiment (Fig 6C). Successful glaucoma induction was confirmed after euthanasia by high degree axonal loss and damage within optic nerve cross-sections (Fig 6D). Transplantation of H7 hRGCs (10^6^ cells/eye) was performed following pars plana vitrectomy and Brilliant Blue G-assisted mechanical ILM peeling over the nasal 2/3 of the macula, followed by fluid-air exchange and puddling of a single cell suspension onto the bare macula (Fig 6B). Tissue was analyzed 8 weeks later.

**Figure 6.**
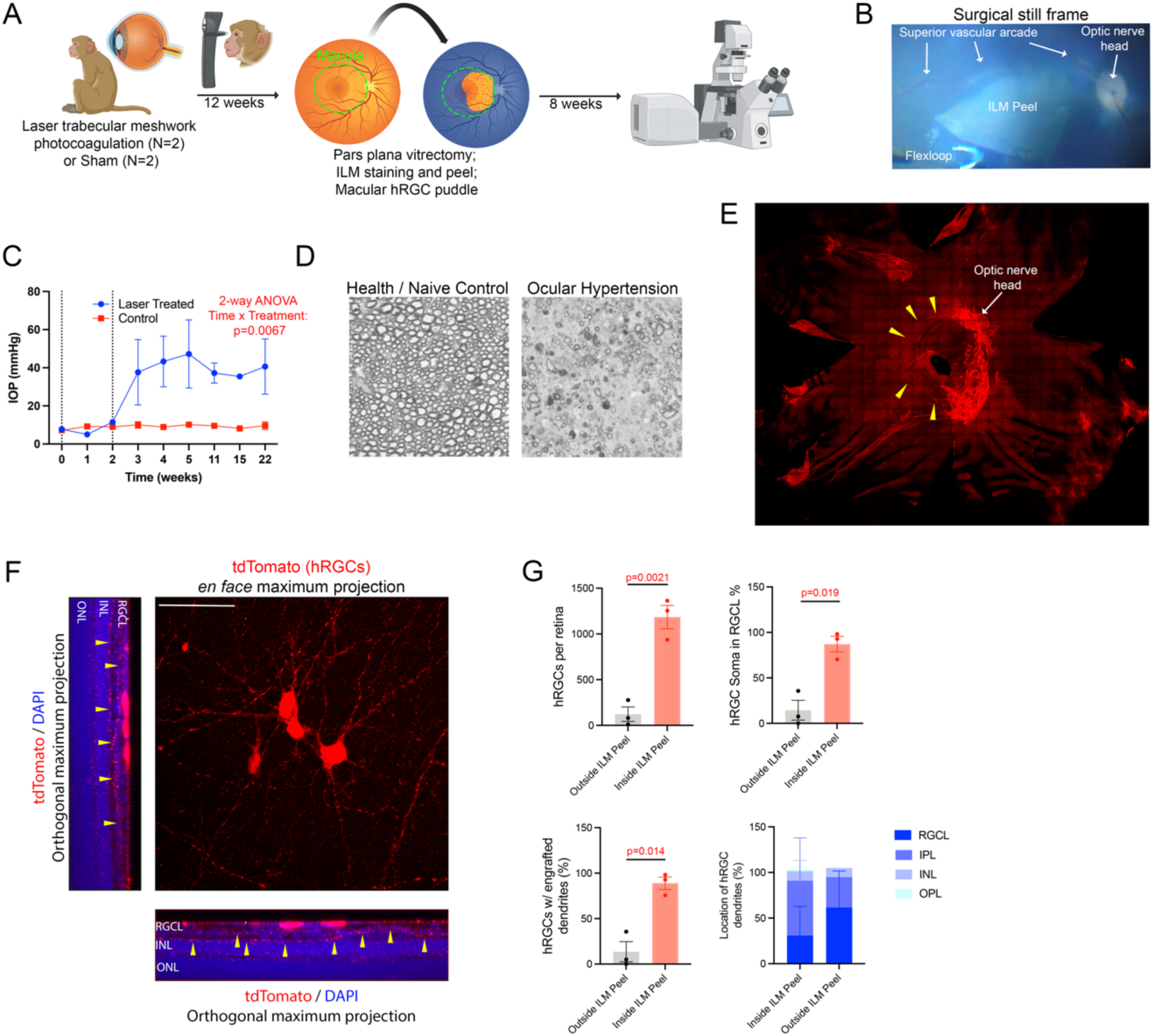
Structural hRGC engraftment in nonhuman primates. Rhesus macaques underwent induction of experimental glaucoma (N=2) or were maintained as healthy controls (N=2). Transplantation of hRGCs occurred following pars plana vitrectomy and ILM peel and histological assessment occurred 8 weeks later. (B) Surgical still frame showing stained internal limiting membrane (ILM) surrounding a zone where the ILM was peeled. (C) Intraocular pressure in animals that underwent unilateral experimental glaucoma induction. (D) Optic nerve cross-sections of healthy eyes and those with experimental glaucoma (ocular hypertension). (E) Retinal flat-mount demonstrating hRGC survival and localization 8 weeks after transplantation. Arrowheads, hRGC axons within the endogenous nerve fiber layer. (F) *En face* and orthogonal projections of confocal z-stack showing engrafted hRGCs. Arrows, dendrites within the inner plexiform layer. Scalebar, 30µm. (G) Quantification of hRGC survival and engraftment at 8 weeks following transplantation. Lower right panel depicts the proportion of surviving hRGC in each retinal layer.

On average, 1,184 ± 221 RGCs survived in 3 of the 4 animals. One retina from a non-glaucomatous animal was subject to significant iatrogenic retinal damage during attempted peeling of a particularly adherent ILM, which was aborted without significant ILM removed. No surviving hRGCs were detected in this retina. Of the 3 remaining animals with surviving hRGCs, >9-fold more hRGCs were localized within the ILM peel bed than in regions of intact ILM (Fig 6G), even though the single cell pool of hRGC cells was applied over the entire posterior retina.

Surviving hRGCs grew extensive neurites including axons within the microarchitecture of the endogenous RNFL superior and inferior arcuate bundles. A subset of these axons navigated towards the ONH (Fig 6E), demonstrating the potential for successful integration of transplanted cells into the complex neural fiber network of the retina. Next, we evaluated the localization of hRGCs somas and dendrites by assessing high-resolution confocal z-stacks of flat-mounted retinas (Supp Video S3; https://youtu.be/PBMKvD2Q0gA). The local percentage of surviving hRGCs for which somas integrated into the RGCL and dendrites extended into the IPL was markedly greater within the ILM peel area compared to areas with intact ILM (Fig 6G).

We also observed tdT labeling of endogenous Muller glia, suggestive of intercellular material transfer from graft-to-host, as we identified previously in mice (Supp Fig S6, Supp Video S4; https://youtu.be/-UHhWHMmzq0) (*40*). Importantly, we did not identify any instances of tdT material transfer from donor-to-host RGCs based on morphology or species-specific antigen labeling.

### Patterns of donor hRGC dendrite and axonal outgrowth

Having observed hRGC dendrite localization to the retinal parenchyma preferentially following ILM disruption (Fig 7A), we next characterized the dendritic morphology of structurally integrated hRGCs in mice. We began by tracing hRGC dendrites within 3D renderings of confocal z-stacks (Fig 7B) and used volumetric analyses to quantify the percentage and length of dendrites within the discrete major retinal layers. Approximately half of hRGC dendrites localized within or just deep to the RGCL, while the remaining half localized to the IPL, as is their native position (Fig 7C). Unlike in organotypic retinal explants, very few dendrites grew ectopically into the inner nuclear layer (INL) and even fewer were detected in the outer nuclear layer (ONL) (Fig 7C).

**Figure 7.**
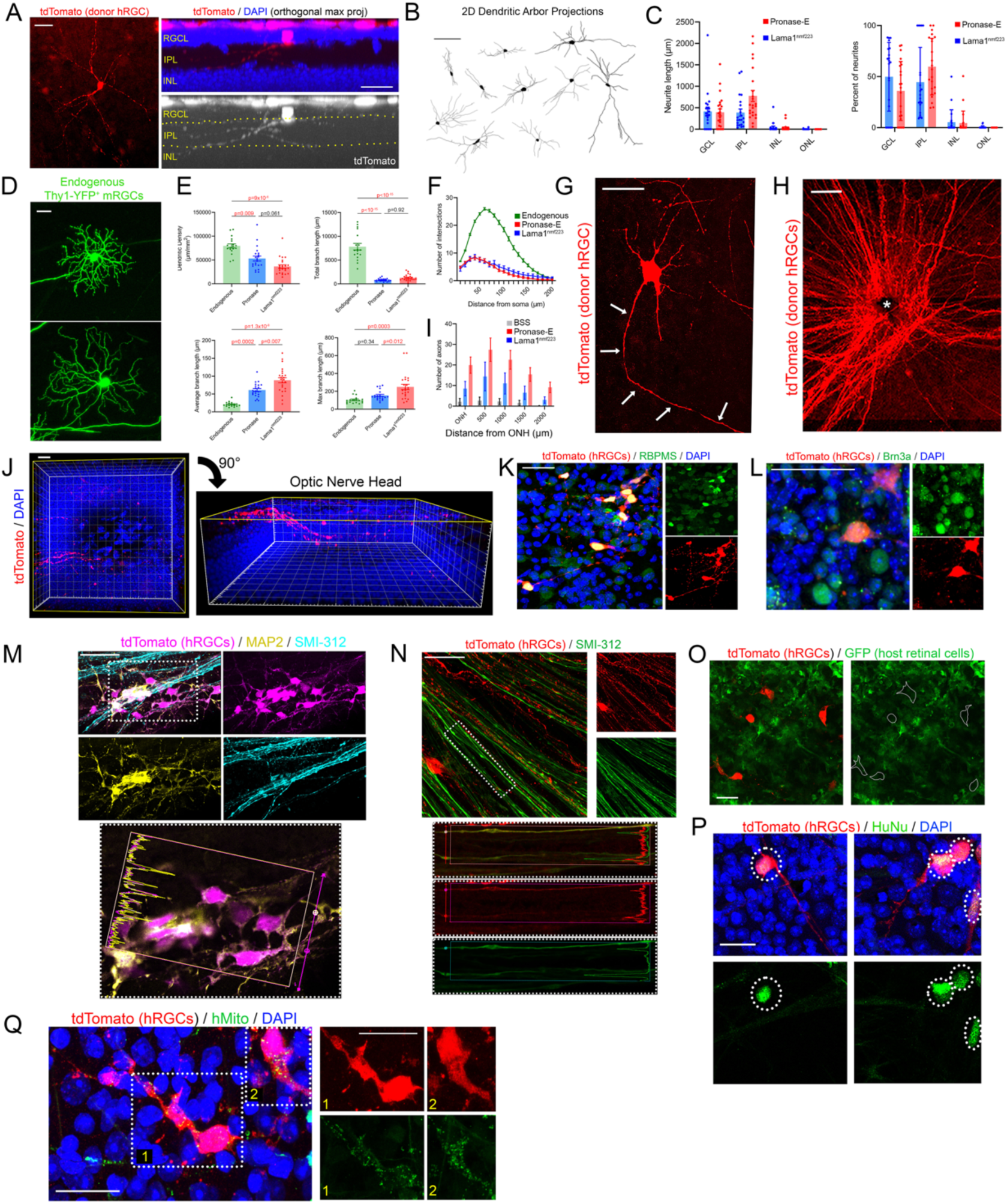
Evaluation of engrafted hRGC dendritic architecture and maturation in mouse retinas. (A) *En face* and orthogonal projections of an engrafted hRGC. Dotted lines delineate boundaries of inner plexiform layer. Scalebar, 20µm. (B) Examples of hRGC dendrite tracings. Scalebar, 100µm. (C) Quantification of hRGC dendrite length and proportion, stratified by localization to each recipient retinal layer. (D) Examples of endogenous murine RGCs in Thy1-YFP. Scalebar, 30 µm. (E) Quantification of dendritic arbor complexity in transplanted hRGCs. (F) Sholl analysis of transplanted hRGC dendritic arbors. (I) Quantification of hRGC axonal growth towards the optic nerve head (ONH) stratified by distance from the ONH. (G) Example of a donor hRGC showing a simple dendritic arbor and one lengthy axon (arrows). Scalebar, 50µm. (H) Maximal projection showing hRGC axons growing into the ONH (*). Scalebar, 100µm. (J) 3D rendering of a confocal z-stack centered at the ONH shows hRGC axons (red) diving deeply into the intraocular ONH. Scalebar, 30µm. Immunofluorescence (IF) for RBPMS (K) and Brn3a (L); Scalebars, 50µm. (L). (M) Immunofluorescence for MAP2 and SMI-312 with linear intensity analysis demonstrating MAP2 expression in hRGC dendrites. Scalebar, 50µm. (N) IF for SMI-312 with linear intensity analysis showing expression in hRGC axons. Scalebar, 50µm. IF for human nuclear antigen (HuNu, P) and human mitochondrial antigen (hMito, Q); Scalebars, 25µm.

We then assessed hRGC dendritic arbor complexity compared to that of mature endogenous mouse RGCs in separate naïve retinas. To do so, we utilized a previously published dataset of confocal micrographs of sparsely labeled RGCs in Thy1-YFP mouse retinas (*59*) (Fig 7D). Compared to endogenous mouse RGCs, integrated hRGCs had reduced dendrite density and total branch length, but greater average branch length and maximal branch length (Fig 7E). This is consistent with an overall lower dendritic complexity with fewer branch points and more unbranched dendrites. Indeed, Sholl analysis demonstrated that the peak dendritic density occurred closer to the soma in donor hRGCs, with a substantially lower overall dendritic complexity (Fig 7F). Together, these data confirm the relatively immature morphology of donor hRGC dendritic arbors at two weeks post-transplantation in mice and provide a benchmark for assessing the maturation of hRGC dendritic architecture over time.

In addition to numerous branched dendrites elaborated for a maximal distance of approximately 200 µm from the soma, hRGCs frequently extended a single lengthy axon that traveled, in some cases, for 2-3mm within the retina (Fig 7G). Remarkably, most of these axons were oriented radially towards the ONH (Fig 2C, 7H). Disruption of the ILM was associated with a significant increase in the number of hRGC axons growing toward the ONH (Fig 7H). hRGC axons descended into the ONH, following the trajectory of the endogenous RNFL as axons enter the optic nerve (Fig 7J, Supp Video S5; https://youtu.be/XGp8FGzaWSo). The retrobulbar optic nerve, however, failed to demonstrate hRGC axons within the nerve parenchyma on histological examination (data not shown), which suggests that either axons face an obstacle at the level of the glial lamina blocking their entry into the nerve or that optic nerve entry simply does not occur by 2 weeks post-transplantation.

### Maturation of hRGCs following transplantation

Differentiation of PSCs to a RGC phenotype yields a mixed population of cells with relatively immature transcriptional signature, for instance, showing only minimal expression of the mature RGC marker RBPMS (*13, 60*). Transplantation of RGCs is associated with molecular evidence of maturation *in vivo* (*9*), so we assessed the molecular phenotype of hRGCs after transplantation. At 2 weeks following transplantation, donor hRGCs uniformly expressed RBPMS, with stronger immunoreactivity in donor neurons than endogenous murine RGCs (Fig 7K). Donor hRGCs also expressed the RGC-specific transcription factor Brn3a (Fig 7L). These data confirm post-transplantation maturation of the donor hRGCs *in vivo*.

We previously noted that hRGCs transplanted onto mouse organotypic retinal explants extended immature neurites which co-expressed dendritic and axonal markers >90% of the time when not engrafted into the tissue. By contrast, almost 25% of hRGC neurites which had engrafted into the retinal parenchyma specified neurite identity, expressing either dendritic or axonal markers, but not both (*2*). Thus, we determined whether neurite maturation occurred in hRGCs following transplantation *in vivo*. As noted above, many hRGCs extended a single long, linear neurite directed towards the ONH which resembled an axon, as well as several shorter more branched neurites which resembled dendrites. Immunofluorescent triple labeling revealed that the shorter neurites near the cell soma expressed MAP2 (a marker of mature dendrites) but not SMI-312 (a marker of mature axons, Fig 7M) whereas the lengthy neurites running with the RNFL expressed SMI-312 (Fig 7N), but not MAP2. Therefore, donor hRGCs transplanted *in vivo* specify mature dendrites and axons within 2 weeks.

### Assessments for graft-to-host intercellular material transfer

To assess the possibility of intercellular material transfer, a phenomenon first described in photoreceptor transplantation studies, whereby donor-derived fluorescent markers are transferred to host neurons and potentially lead to erroneous assumptions of donor cell engraftment (*61–65*). we first transplanted tdT^+^ hRGCs into mice ubiquitously expressing green fluorescent protein (GFP) under the CAG promoter. All putative hRGCs which were tdT+ were GFP-negative, confirming the donor origin (Fig 7O). Further, we immunolabeled retinas from mice that received hRGC transplants with antibodies that specifically recognize human nuclei (HuNu, Fig 7P) or human mitochondria (hMito, Fig 7Q) and noted subcellular immunoreactivity in all purported donor hRGCs, again confirming their donor origin. The localization of hMito to hRGC neurites (Fig 7Q) enabled clear identification of donor dendrites and axons, in addition to cell bodies.

### Functional retinal integration of donor hRGCs

Finally, we sought to determine whether donor hRGCs formed synapses with host inner retinal neurons following transplantation in mice. We immunolabeled flat-mounted retinas following transplantation with antibodies recognizing presynaptic synaptophysin and postsynaptic PSD-95 and noted extensive colocalization of punctate synaptic machinery within the dendrites of donor hRGCs (Fig 8A).

**Figure 8.**
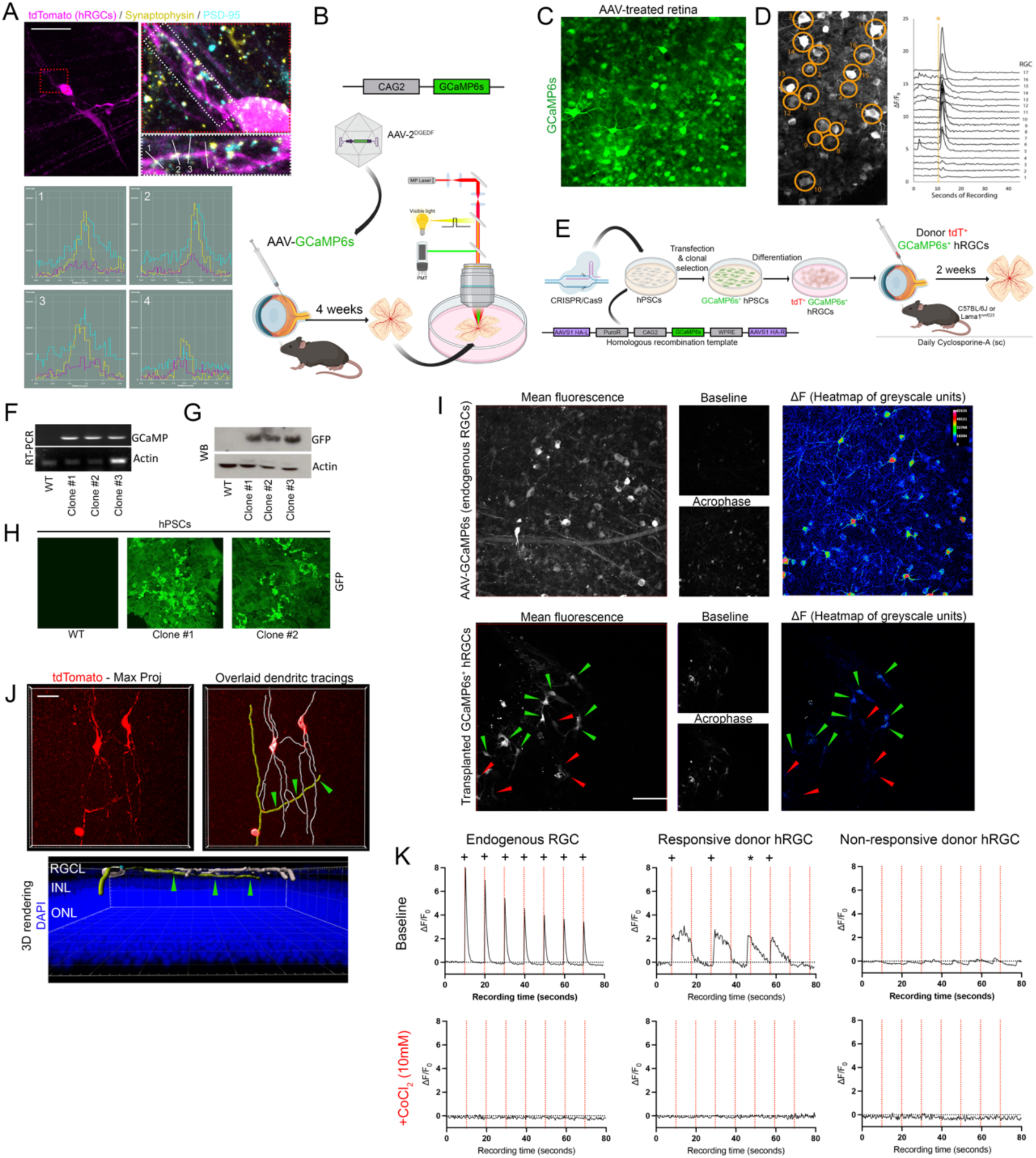
Functional engraftment of hRGCs in mice. (A) Immunofluorescence for presynaptic synaptophysin and post-synaptic density (PSD)-95 with linear intensity analyses demonstrating co-localization at hRGC dendrites in the inner plexiform layer. (B) C57BL/6J mice underwent intravitreal injection of AAV driving expression of GCaMP6s, followed 4 weeks later by 2-photon calcium imaging with light-evoked retinal simulation. (C) Fluorescent GCaMP6s in AAV-transduced endogenous RGCs. (D) Example of calcium imaging in AAV-transduced endogenous RGCs. Still frame is shown in the left panel where individual RGCs are circled and labeled. Plot of ΔF/F_0_ as a function of time shown in the right panel for each of the labeled RGCs. Visible light stimulation occurred at 10sec of recording (*orange line). (E) H7 hESCs were gene edited to knock GCaMP6s into the AAVS1 safe harbor locus, and hRGCs were differentiated from this line and transplanted into mice for subsequent calcium imaging 2 weeks later. RT-PCR (F), Western blot (G), and immunofluorescent validation (H) of GCaMP6s in knock-in PSC lines. (I) Exemplar calcium imaging still frames from endogenous RGCs (top row) and transplanted hRGCs (bottom row). Baseline images are unstimulated and acrophase images are from peak GCaMP6s fluorescence. Heatmaps show the difference in fluorescent intensity from baseline to acrophase. Green arrowheads, light responsive hRGCs. Red arrowheads, non-light responsive hRGCs. (J) Following calcium imaging, retinal flat-mounts were processed for confocal microscopy. 3D reconstructions demonstrated that light-responsive RGCs elaborated dendrites in the inner plexiform layer (yellow dendrite tracing, green arrows) while those that were non-light responsive had dendrites that remained superficial to the retina. Scalebars, 30µm. (K) Plots of ΔF/F_0_ as a function of time for endogenous RGCs and both light-responsive and non-responsive transplanted hRGCs before and after exposure to CoCl_2_ (10mM). Visible light stimulation was given every 10sec (red lines). +, light-evoked calcium response. *, no response following a spontaneous calcium transient that preceded stimulation.

We next leveraged optical electrophysiology with a genetically encoded calcium indicator (GCaMP6s) composed of GFP, calmodulin and the M13 peptide sequence from myosin light chain kinase to assay intracellular calcium fluxes associated with action potential firing (*66*). In RGCs, photoreceptor light stimulation is followed by synaptic transmission to bipolar cells and then to RGCs, resulting in a transient train of RGC action potentials that is associated with a short burst in intracellular calcium (*67, 68*). Thus, RGC GCaMP6s expression should yield a transient increase in fluorescence coincident with a burst of action potentials, enabling visualization in real-time of their light-responsive electrophysiologic activity. We confirmed this light response activity using two-photon microscopy of *ex vivo* mouse retina preparations transduced 4 weeks earlier with a variant AAV vector (DGEDF) that was developed through directed *in vivo* evolution to yield strong expression in inner retinal neurons following intravitreal injection (*69*), and which drove GCaMP6s expression under control of the CAG2 promoter (Fig8B, C). Consistent, strong calcium fluxes were observed across most, but not all, endogenous RGCs using time-lapse imaging of GCaMP6s in conjunction with full-field visible light stimulation (Fig 8D, I, Supp Video S6; https://youtu.be/hmo4LPEAKp0). The amplitude of response decreased over consecutive stimulations (Fig 8K, Supp Videos S7-8; https://youtu.be/28dghXrnjQ4 and https://youtu.be/ZSvnJYftf6I). Variable responsivity is likely explained by the feature selectivity of diverse RGC subtypes, as orientation and direction-selective RGCs would not be expected to respond to a full-field light stimulus.

We then generated ubiquitously expressing PSC lines using CRISPR/Cas9-mediated gene editing to knock GCaMP6s into the AAVS1 safe harbor locus (Fig 8EF-H). *In vivo* transplanted GCaMP6s^+^ hRGCs frequently exhibited spontaneous calcium transients and demonstrated a noisier baseline than endogenous mouse RGCs (Fig 8I, K). A subset of hRGCs responded to light stimuli, though responses were generally weaker and longer in duration than for endogenous mouse RGCs and many hRGCs did not respond to every stimulation within a series (Fig 8I, K, Supplemental Videos S9-12; https://youtu.be/cXxSzp1j-Sk, https://youtu.be/caFTSvs4rFY, https://youtu.be/i2WOObEQ0QM, and https://youtu.be/aSYA2XC-wjo). hRGCs in Lama^1nmf223^ retinas were significantly more likely to exhibit light responsivity (49 of 273 hRGCs (18%) across 5 retinas) than when transplanted into C57BL/6J eyes with intact ILM (3 of 32 hRGCs (9%) across 5 retinas), confirming that ILM disruption promotes the integration of transplanted RGCs. To ensure that light responsivity was mediated through synaptic integration with endogenous retinal circuits and not through intrinsic photosensitivity, we added cobalt chloride (CoCl2, 10mM) to the bath to block synaptic transmission (*70, 71*) and observed complete loss of light-evoked calcium fluxes in endogenous RGCs of AAV^DGEDF^-GCaMP6s retinas and in GCaMP6s-hRGCs within transplanted retinas (Fig 8K). Correlational confocal microscopy of fixed retinas confirmed that light-responsive hRGCs extended dendrites into the IPL. Collectively, these data demonstrate that ILM disruption not only enables structural integration of hRGCs into the IPL but also enables synaptic integration into functional inner retinal circuits.

## DISCUSSION

Transplantation of human PSC-derived RGCs holds potential for restoring vision across an array of optic neuropathies. However, several critical obstacles hinder the efficacy of this approach for restoring visual pathway function, including limited survival and functional engraftment of neurons transplanted into the vitreous cavity. Here, we identify and overcome the ILM as a primary barrier to the engraftment of intravitreally transplanted RGCs. We demonstrate that the ILM prevents most donor hRGCs from migrating into the recipient inner retina and from elaborating dendrites into the IPL, where synaptic input from afferent bipolar and amacrine cells might be established. Using two independent approaches – proteolytic ILM digestion and a developmental model of ILM dysgenesis– we observed significant increases in structural hRGC engraftment into the inner retinas of mice. Moreover, we found that disruption of the ILM in a mouse model of glaucoma, human postmortem retinal tissue, rats, and nonhuman primates similarly augments the engraftment of transplanted hRGCs. Remarkably, many of the structurally engrafted hRGCs were found to gain light sensitivity through a mechanism that was dependent on synaptic transmission, suggesting functional integration into inner retinal circuits. Together these data provide solutions to some initial roadblocks for future work that is aimed at replacing functional RGCs within the degenerated anterior visual pathway.

By providing a clearly translatable avenue for increasing donor hRGC engraftment into the inner retina, we hope to bring the possibility of transplantation-mediated RGC replacement closer to clinical application. Indeed, ILM peeling performed after pars plana vitrectomy is a widely implemented surgical technique for treatment of the vitreomacular traction, macular hole, and epiretinal membrane in older persons (*72, 73*). However, it is possible that complete local ILM removal may introduce new challenges to hRGC engraftment. Neurodevelopmental studies have established that signaling between RGCs and the ILM is critical for organized patterning of the RGC layer and dendritic projections to the IPL. β1-integrin recognition of ILM is required for RGC organization into a single-celled layer. In the absence of β1-integrin, or Cas adaptor proteins which signal downstream of integrins, retinal neurons form ectopic cell agglomerations, extravasate past the ILM, and in some ways phenotypically resemble Lama1^nmf223^ mutants (*74*). In addition, loss of laminin-γ3 causes defects in retinal lamination and differentiation of RGCs. As such, cellular permeabilization of the ILM, sub-ILM injections (*75*) laser photoporation (*76*), or surgical repositing of the ILM over the donor hRGC graft may be more advantageous for RGC engraftment than *en bloc* ILM resection. Regardless, the barrier role of the ILM must be accounted for in RGC transplantation.

In addition to its translational relevance, our findings provide a new avenue for experimental research on neuroplasticity and circuit development. Numerous molecular pathways have been reported to dramatically enhance the axon regeneration of injured endogenous RGCs through the optic nerve, past the chiasm, and to central subcortical visual targets (*77*). However, experiments aimed at connecting donor RGCs to inner retinal neurons and growing their axons into the optic nerve have been hampered by extremely poor engraftment into the retinal parenchyma (*7*). We have now established a benchmark for hRGC dendritic lamination within the IPL to evaluate additional manipulations that enhance the effectiveness of this process. Moreover, given the localization of synaptic machinery to hRGC dendrites within the IPL and functional integration into light-sensitive circuits, robust functional assessments of donor neuron connectivity to afferent neurons can now be explored. Finally, given the spontaneous growth of hRGC axons towards and into the ONH, the development of methods that enhance the extension of axons into the retrobulbar optic nerve and beyond are now within reach. The directed growth of hRGC neurites towards the ONH in rodents and non-human primates is intriguing and suggests either that hRGC axons use the existing RNFL as a conduit (potentially mediated by cell surface receptor interactions) or that yet-unidentified axonal guidance cues are present in the adult retina.

Despite the advances documented here, hurdles to efficient RGC replacement remain. Low survival of transplanted hRGCs is a pervasive issue and must be improved to yield a viable regenerative therapy. Though ILM disruption increased hRGC survival in most of the models we evaluated, absolute survival remained suboptimal. At least three likely contributors to donor hRGC cell loss following transplantation should be addressed in future work. These include cell-intrinsic induction of apoptosis in this relatively delicate cell type, immunological insult to the graft, and lack of appropriate microenvironmental support for the neurons.

Neurons are dependent on a highly regulated extracellular microenvironment, which is established *in vivo* by supportive glia, but must be recapitulated experimentally in transplantation paradigms. RGCs are vulnerable to injury of subcellular compartments including dendrites and axons, which are likely to be damaged during purification and preparation for transplantation. Indeed, detachment of cells from growth substrates, or *anoikis*, triggers caspase-8 dependent apoptosis (*78*) and may contribute to early cell death following transplantation. Neuroprotective strategies to prevent RGC death from a range of insults have identified numerous intrinsic cell signaling cascades that prevent RGC death from applied insults *in vitro* and from optic neuropathy *in vivo* (*79–81*). Supportive neuroprotection to RGC transplants may yield improved survival rates.

Cell transplants are subject to both innate and adaptive immune activity which undermine survival and engraftment. The current study demonstrated that immunosuppression with cyclosporine yielded graft survival that was commensurate with that achieved by their transplantation into NSG mice, suggesting that immunological graft rejection (i.e., T cell-mediated immunity) is adequately controlled by immunosuppression over the first two weeks following transplantation. It is likely that innate neuroinflammatory activity plays an additional role in triggering hRGC death early after transplantation. Reactive microgliosis and astrogliosis are known to propagate neurotoxicity in several neurodegenerative conditions including glaucoma (*82, 83*). Pharmacologic approaches to ameliorating the micro- and macroglial responses to optic neuropathy and RGC transplantation may support enhanced graft survival (*84*).

Finally, augmentation of the recipient microenvironment may improve hRGC survival at early time points. The vitreous cavity, where we transplanted hRGCs in this study, is relatively hypoxic and devoid of direct metabolic support for neurons. It is possible that the hRGCs that survive transplantation do so owing to their ability to migrate into the retina where they can obtain metabolic support derived from retinal vascular perfusion. Thus, increasing the early engraftment of donor hRGCs into the host retina may be critical for ensuring survival. This theory is substantiated by our observation that ILM disruption enhanced survival in most of the models tested. In addition, RGCs are dependent on neurotrophin support, especially during development. Brain-derived neurotrophic factor (BDNF) transported retrogradely from postsynaptic subcortical targets rescues RGCs from developmental apoptosis (*85*). BDNF supplementation is neuroprotective to endogenous RGCs following injury (*86, 87*), and delivery of sustained release BDNF (along with CNTF and GDNF) appears to promote hRGC survival following transplantation (*88*). Thus, co-delivery of hRGCs with neurotrophins may, in future studies, improve graft survival at early time points when the neurons are most susceptible to death.

The present experiments extend findings from previous reports in the regenerative medicine field. hESC-derived RGCs have been transplanted into rat eyes, where somal co-localization with the RGCL was reported. However, donor cells did not appear to extend neurites by 3 weeks post-transplantation (*89*). In addition, those donor neurons did not express RBPMS, leaving open the question of whether the donor cells had matured into a functional RGC phenotype. Vrathasha et al. (*34*) transplanted AAV-GFP labeled human iPSC-derived RGCs into immunocompetent mice and reported a remarkable degree of survival, dendritic complexity, and light responsivity at 2-5 months following transplantation. Vanugopalan et al. (*31*) demonstrated that a small proportion of primary mouse RGCs transplanted into rat recipients with intact ILM appear to survive, attain mature dendritic morphology resembling endogenous RGCs, achieve synaptogenesis, and conduct electrophysiological responses to light stimulation, despite overall low survival rates. Differences in outcomes might be explained by disparities in the donor cell source or preparation, or the recipient model. However, in comparison to those prior studies, a notable strength of the current data is inclusion of rigorous control experiments that ruled out the possibility of intercellular material transfer, which could lead to erroneously identifying endogenous murine RGCs as donor-derived neurons (*90*). In our view, this is the first demonstration of functional engraftment by transplanted RGCs that are definitively donor-derived.

We rigorously analyzed the localization of hRGCs within the host retina, using flat-mounted retinas rendered and analyzed in 3D. This approach enabled us to simultaneously evaluate the full extent of hRGC dendritic arbors in the x,y plane while retaining the ability to resolve z positions within specific retinal layers. Critically, previously reports have focused exclusively on analyses of retinal sections, which capture only a fragment of horizontally oriented dendritic arbors, or 2D maximal projection images of retinal flat-mounts, which provide no discrimination between cells lying on the retinal surface versus those which may have migrated below the ILM (*9, 31, 33, 91, 92*). Studies in pre-clinical models of RGC injury suggest dendritic extension into the RGCL by donor neurons, which was described as neuronal engraftment even though this dendritic position would not enable synaptogenesis with bipolar and/or amacrine cells (*33*). Indeed, methods to determine graft localization and engraftment are inconsistent across published literature, and some of the above studies even suggest that transplanted RGCs that accumulated on the vitreous side of the ILM are integrated. These observations are inconsistent with the native or desired localization of RGCs and point to the need for more standardized methodologies of determining cellular localization (*7*). In our view, it is insufficient to assess functional integration into the host retinal circuitry without first demonstrating dendrite distribution into the retinal IPL (*33*). The lack of consistent methodology for discerning donor somal coplanarity with the host RGCL, and assessment of integration into the IPL, limits accurate reporting and comparison of transplantation outcomes. We propose that the approach taken here is one solution to this issue.

An alternative to intravitreal transplantation of hRGCs has been proposed by Soucy et al. (*88*) that involves subretinal transplantation of the neuronal graft. By circumventing the ILM altogether, a small number of RGCs can migrate into the inner retina from the subretinal space. Delivering a guidance signal, such as recombinant stromal cell-derived factor-1 (SDF-1), into the vitreous cavity further enhances the migration of hRGCs towards the RGC layer. However, the ultimate efficiency of this delivery method remains to be determined and would be technically more demanding and riskier than intravitreal transplantation. Moreover, it will be important to demonstrate that the migrating neurons do not disrupt the neurocircuitry of the recipient retina.

The specific techniques utilized here to bypass the ILM come with important considerations. The developmental ILM dysgenesis model is based on a mutant mouse and is not readily translatable to other species. The use of pronase-E to enzymatically digest the ILM also comes with caveats. As noted in the *Methods*, we have found that the manufacturer-reported enzymatic activity of pronase does not correlate well to its observed effects in the eye, necessitating that end users either acquire the same lots reported here or perform a lot-specific titration bioassay to determine the optimal concentration of pronase that will digest the ILM without causing intraocular bleeding, neurotoxicity, or gliotoxicity. It is also worth noting that since the enzymatic activity (U/mg) varies across lots of enzyme, so reporting the dose of enzyme administered in percentage (g/100mL) is insufficient (*61*) and instead should be reported in terms of enzymatic units based on lot-specific conversion of enzymatic activity per mg of enzyme. We have documented destructive effects with high doses of intraocular pronase (Fig 1B), and it is likely that the concentration drives the differences between the effects of pronase noted here and elsewhere. As an additional potential challenge, sensitivity to pronase digestion may vary patient to patient.

In conclusion, we demonstrated that the ILM is a key barrier to the engraftment of human stem cell derived RGCs into the host mammalian retina, and that ILM disruption enables a significant proportion of donor RGCs to extend dendrites into the IPL where they receive presynaptic visual input for eventual communication to the brain. Future research aimed at increasing donor hRGC survival post transplantation, evaluating and modulating hRGC integration into specific retinal circuits, and driving hRGC axonal connectivity to the brain will likely benefit from ILM disruption as a primary enabling step in the transplantation process.

## MATERIALS AND METHODS

### Animals

Mice, rats, and nonhuman primates were housed on a 12h light/dark cycle with food and water available *ad libitum*. *See Supplemental Methods (SM) for details*. All animals were treated in accordance with the ARVO Statement for the Use of Animals in Ophthalmic and Vision Research and according to experimental protocols approved by the Johns Hopkins Animal Care and Use Committee.

### Human Tissue

Postmortem human donor eyes were obtained from a 72-year-old male and an 87-year-old female from the Lions Gift of Sight Eyebank (St. Paul, MN) and had no known history of ocular pathology. Globes were shipped on ice and processed with 24h of death.

### Organotypic Retinal Explants

Human retinas were separated from the RPE and a 5mm trephine used to obtain retinal discs from the macula. Peripheral retina and foveal tissue were not evaluated. Mouse retinas were isolated immediately following euthanasia and dissected from the RPE. Four relaxing incisions were placed to flat-mount entire murine retinas in Maltese-cross configuration. Both human and mouse retinal explants were cultured on polytetrafluoroethylene organotypic filters (Millipore-Sigma) with the photoreceptor side down, at an air/medium interface maintained in humidified incubators with 5% CO_2_ at 37°C. Media (iNS) consisted of Neurobasal-A supplemented with 1% N2, 2% B27, 2mM GlutaMAX, 100U/mL penicillin, 100µg/mL streptomycin (*59, 93*). A 2µL droplet of 1.5×10^4^ hRGCs was transferred onto the surface of each explant 24h after dissection.

### Enzymatic ILM Disruption

Pronase-E (Millipore-Sigma, Burlington, USA, Cat #P8811) was injected intravitreally (mice: 1µL; rats: 5µL) at the indicated concentrations. Atop human retinal explants, Pronase-E (3mU in 5µL) was applied. The enzymatic reaction was terminated after 1h with 300μL of inhibitor solution (0.15 mM BSA and 0.375 mM Ovomucoid in BSS), followed by washout with 3 mL of BSS. *See SM for details*.

### Immunosuppression

Cyclosporine was administered to rodents by daily subcutaneous injection (mice: 25mg/kg/day; rats: 10mg/kg/day). Rhesus macaques received methotrexate (5mg/kg/week intramuscular) and cyclosporine (15mg/kg/day subcutaneous, targeting a whole blood trough concentration of 150-250ng/mL) starting 1 month before transplantation. Methylprednisolone was administered at 1mg/kg/day for one week following transplantation and then converted to methylprednisolone acetate intramuscular injections (22.4mg/kg/week).

### hRGC Differentiation and purification

Differentiation of human PSCs containing the *BRN3B-tdT-mThy1.2* was achieved as described previously (*13*). *See SM for details*.

### GCaMP6s^+^ hPSC line generation

H7 hESCs containing the *BRN3B-tdT-mThy1.2* construct were subject to CRISPR/Cas9-mediated knock-in of GCaMP6s under control of the CAG promoter at the AAVS1 safe harbor locus. See SM for details.

### GCaMP6s Adeno-Associated Virus

To develop methodology for light-evoked multiphoton calcium imaging of RGCs in situ, AAV-2^DGEDF^ (*69*) encoding GCaMP6s was injected intravitreally in mice (1µL, 2.5×10^13^vg/mL) 4 weeks prior to imaging. *See SM for details*.

### Intravitreal injection

Mice were anesthetized (inhalational isoflurane 2.5%) and pupils dilated with 2.5% phenylephrine hydrochloride (Akorn, Inc). Ophthalmic gel (2.5% Goniosoft, OCuSOFT, Rosenberg, USA) was used to couple a 4mm glass contact lens to the cornea and enable direct visualization of the fundus and needle tip, to avoid lens penetration/contact or damage to the retina during intravitreal injection. A 10µL Hamilton syringe and 33-gauge metal needle (Hamilton Co., Reno, USA) were used to deliver payloads through transretinal injection 3mm posterior to the superotemporal limbus. The needle was maintained in the vitreous cavity for 1min after injection to allow equilibration of the IOP and diffusion of the injected solution while minimizing reflux.

### Mouse ocular hypertensive glaucoma

Intracameral microbead injection was used to induce ocular hypertension in mice, as described previously (*94*) and modified from Sappington et al. (*95*). *See SM for details*.

### Non-human primate ocular hypertensive glaucoma

Experimental glaucoma was induced using a protocol derived from Dr. Brad Fortune (Devers Eye Institute) (*96–98*). Rhesus macaques were sedated with ketamine and dexmedetomidine. IOP was measured using an iCare IC200 (iCare USA). Proparacaine hydrochloride 0.5% (Bausch & Lomb) was applied and a Kaufman laser lens was placed on cornea coupled with Goniosoft. Two animals underwent laser photocoagulation of the trabecular meshwork (TM) unilaterally using a slit lamp equipped with 532nm diode laser. Laser spots were placed in confluent fashion for 270 degrees (laser power, 360-400mW; spot size = 125µm; 1sec/spot) until good blanching of the trabecular meshwork and some spillover blanching of the ciliary body were observed. Two laser sessions spaced 2 weeks apart were required to achieve IOP elevation. The IOP was measured weekly for a month and then monthly until euthanasia.

### Non-human primate hRGC transplantation

Pars plana vitrectomy occurred under general anesthesia with animals positioned supine. The operative eye was dilated, a pediatric lid speculum was placed, and three 25G trocars were inserted 2.5mm posterior to the limbus. Core vitrectomy was performed, intravitreal triamcinolone acetate was instilled to visualize the vitreous, and the posterior hyaloid was detached using a cutter. Brilliant Blue-G was instilled to stain the ILM and a flexloop was used to create an ILM edge. ILM forceps was used to peel the ILM from the optic disc to just temporal to the fovea over approximately 2/3 of the macula. Air-fluid exchange was performed and 10^6^ hRGCs in a single cell suspension containing iNS media were injected over the posterior pole. Air was then exchanged for SF_6_ gas and subtenon Kenalog was administered. The eye was left facing upward for 45min to facilitate cell attachment to the retina, prior to surgical closure and recovery.

### Confocal Microscopy

Retinal flat-mounts were examined using a LSM Zeiss 880 confocal microscope (Carl Zeiss Microscopy, Thornwood, USA). For cell quantification and topographic analyses, wide-field full-thickness confocal microscopic z-stacks were captured using a Plan-Apochromat 10X/0.45 M27 objective and image tiles were stitched. For high magnification analysis of individual or groups of cells, full-thickness confocal microscopic z-stacks were obtained using a C-Apochromat 40X/1.2 W Korr UV-VIS-IR. The pinhole was set to 1 Airy unit. Scans were obtained in line sequential mode at a pixel dwell time of 1.54µs using bidirectional scanning. Excitation detection wavelengths 488 and 633 shared a single track, whereas excitation wavelengths 405 and 561 had individual tracks to avoid signal bleed-through. Microscopists were masked to treatment groups to ensure unbiased field selection for imaging.

### Quantification of neuronal survival

Quantification of cell survival on retinal flat-mounts was performed using the ImageJ (National institute of health, USA) cell counter plugin (*99*). NeuN^+^ bipolar and amacrine cells in the RGCL and PKCα^+^ rod bipolar cells in the inner nuclear layer were quantified in four quadrants per retina at an eccentricity approximately halfway between the ONH and the retinal edge. Donor tdT^+^ hRGCs were quantified manually to minimize error, scrolling plane by plane through confocal z-stacks to avoid misinterpretation. All surviving hRGCs were quantified within each retinal flat-mount, rather than sampling and extrapolating overall donor neuron survival.

### Assessment of donor hRGC soma and neurite integration

hRGC soma locations were determined to be superficial to, within, or deep to the recipient RGCL using high magnification confocal z-stacks and Zen software (v8.1.0, 2012 SP1, Carl Zeiss Microscopy), as previously described (*2, 100*). Stack scrolling and orthogonal projections were used to visualize donor hRGC somas and the host RGCL concomitantly, and hRGCs were designated as coplanar with the RGCL only if there was not an endogenous RGC soma deep to the hRGC. Investigators were masked to the identity of treatment groups during image analysis.

Analyses of hRGC neurite localization within the recipient retina were conducted using Imaris (v9.3, Oxford Instruments, Zurich, Switzerland) as previously described (*2, 100*). High magnification confocal z-stacks were rendered in 3D and individual hRGC neurite processes were traced in semi-automated fashion using the filament and autopath tools. Local DAPI staining of the recipient retinal nuclear layers was used to identify neurite localization. The length of neurites within specific retinal layers was quantified using Imaris by segmenting 3D neuronal tracings according to the DAPI-registered retinal layer. Quantitative metrics describing the dendritic morphology of structurally integrated hRGCs were obtained using Imaris. Investigators were masked to the identity of treatment groups during image analysis.

### Calcium Imaging

After dark adaptation for >6h, retinae were explanted under dim red light onto organotypic membranes with the photoreceptors facing down and maintained in a perfusion chamber running carbogenated iNS media at 35°C and a rate of 1mL/min. Zeiss LSM 710 confocal microscope (Carl Zeiss Microscopy) was used with a 20x W Plan Apochromat objective (NA 1.0). Fluorophore excitation was achieved with a two-photon laser source (Chameleon Ultra Vision II, Coherent, Inc, Santa Clara, CA, USA), tuned to a wavelength of 930nm. Emission was collected through a 500-550 nm bandpass filter at 3 Hz. To stimulate retinal phototransduction, tissue was exposed to visible light stimulation using the fluorescence recovery after photobleaching (FRAP) function with a 405nm laser. Stimuli were presented 5-10 times sequentially at 10sec intervals. To inhibit the synaptic transmission within the retina, 10mM of cobalt chloride was added to the perfusate. Data were analyzed using Suite2P (Howard Hughes Medical Institute Janelia Research Campus) (*101*). After calcium imaging, retinas were then PFA-fixed for 1hr and immunolabeled for standard confocal microscopy.

### Statistical Analyses

Data are reported as mean ± SEM unless otherwise stated. Each experiment was performed at least twice. Group means were compared using unpaired non-parametric Mann-Whitney U tests or non-parametric Kruskal Wallis analyses of variance. Following Bonferroni correction for multiple comparisons, p < 0.05 was considered statistically significant. Data were analyzed and plotted using Prism (8.0, GraphPad Software, San Diego, USA).

## ACKNOWLDGEMENTS

The authors are grateful to Dr. Jessica Izzi, DVM, Dr. Lydia Hopper, PhD, and their teams within the Johns Hopkins Research Animal Resource and Non-human Primate Behavior departments for their assistance with and outstanding care of the rhesus macaques used in this study. The authors are grateful to Mimi Santander, RN and Robin Harrison, CST for their expert assistance with vitreoretinal surgery in non-human primates. The authors gratefully acknowledge Dr. Shannon Boye (University of Florida) for sharing the AAV^DGEDF^ capsid and Dr. Xitiz Chamling (Johns Hopkins University) for sharing the AAVS1 CRISPR/Cas9 targeting plasmid used in this study. Schematic diagrams in figures were created using Biorender.com.

## FUNDING

National Institutes of Health grant K08EY031801 (TVJ)

National Institutes of Health grant R21EY034332 (TVJ)

National Institutes of Health grant P30EY001765 (DZ)

Research to Prevent Blindness Career Development Award (TVJ)

Research to Prevent Blindness unrestricted funding to the Wilmer Eye Institute

Bright Focus Foundation National Glaucoma Research Award G2002005S (TVJ)

The Zenkel Family Foundation Research Award (TVJ)

Department of Defense Focused Translational Team Science Award #13752359 (TVJ and DZ)

The Glaucoma Foundation Rajen Savjani and Joe Rosen Grant Awards (TVJ)

## AUTHOR CONTRIBUTIONS

Conceptualization: TVJ

Methodology: EAA, MMV, KYZ, AN, EK, and TVJ

Investigation: EAA, BB, MMV, SM, KYZ, JD, AN, EK, JTH, and TVJ

Visualization: EAA, BB, MMV, SM, KYZ, JD, AN, WY, EK, and TVJ

Funding Acquisition: DZ and TVJ

Project Administration: JD, EK, and TVJ

Supervision: JD, EK, and TVJ

Writing – original draft: EAA, BB, EK, and TVJ

Writing – review & editing: EAA, BB, MMV, EK, SM, KYZ, JD, AN, WY, SQ, SH, HAQ, DJZ, JTH, and TVJ.

## COMPETING INTERESTS

The authors have no competing interests to report.

## DATA AND MATERIALS AVAILABILITY

Data are available from the corresponding author upon reasonable request. Cell lines, plasmids, and vectors are available from the primary author upon reasonable request and subject to a material transfer agreement.

## SUPPLEMENTAL METHODS

### Animals

Adult (8-12 weeks) male and female C57BL/6J (Jackson strain #664), heterozygous C57BL/6-Tg (CAG-EGFP)1Osb/J (Jackson strain #3291), and B6.Cg-Tg(Thy1-YFP)HJrs/J (Thy1-YFP, Jackson Strain #3782) mice were obtained from Jackson Labs (Bar Harbor, USA). Homozygous C57BL/6J-Lama1^nmf223^/J mice (Jackson strain #5094) were generously donated by the Malia Edwards Laboratory (Johns Hopkins University, JHU). NOD.Cg-*Prkdc^scid^Il2rg^tm1Wjl^/Szj* (NSG, Jackson strain #5557) were obtained from the JHU Research Animal Resources breeding program. Wister rats were obtained from Charles River Laboratories (Wilmington, MA). Rhesus macaques (1 male and 3 females) were 3-4 years old, weighed 4-7kg, and were obtained from the JHU Research Animal Resources breeding Program.

### Enzymatic ILM disruption

Pronase-E, a nonspecific mixture of endo- and exoproteases from *Streptomyces griseus*, was reconstituted in balanced salt solution (BSS) at the indicated concentrations. The manufacturer reported enzymatic activity of pronase is defined such that one unit hydrolyzes casein to produce 1.0 µmole of tyrosine per minute at pH 7.5 and 37°C. However, enzymatic activity depends upon the substrate and the microenvironmental conditions. We found that the intraocular effects of pronase were consistent at specific concentrations of reported enzymatic activity (in U/mL) within given lots, but that similar enzymatic activity concentrations from different lots produced inconsistent effects within the eye. Therefore, rather than rely on manufacturer reported enzymatic activity, we performed a dose-titration bioassay for individual pronase lots (Fig 1).

### Human pluripotent stem cell culture

H7 and H9 hESCs (originally from WiCell Research Institute, Madison, WI), and EP1 iPSCs containing the *BRN3b-tdT-mThy1.2* construct were maintained in mTeSR1 media (Stem Cell Technologies, Vancouver, USA) on growth factor-reduced Matrigel-coated (BD Biosciences, Franklin Lakes, USA) plates at 37°C in 5 % CO2. Stem cells were passaged by dissociation with Accutase (Stem Cell Technologies) and re-plating in mTeSR1 media for maintenance with 5 µM blebbistatin (Millipore-Sigma, Burlington, MA).

### hRGC Differentiation and purification

PSCs were dissociated using Accutase and 5×10^5^ cells were seeded on Matrigel coated six-well plates in mTeSR1 with 5µM blebbistatin (Millipore-Sigma). After 24h, mTeSR1 was completely exchanged to iNS media consisting of 1:1 Neurobasal and DMEM/F12 media supplemented with 1% N2, and 2% B27. Beginning 24h later, fresh iNS media supplemented with small molecules was exchanged every other day. The working concentrations of small molecules were: Forskolin (FSK, 25µM, Stem Cell Technologies). Dorsomorphin (DSM, 1µM, Stem Cell Technologies), Inducer of definitive endoderm 2 (IDE2, 2.5µM, Stem Cell Technologies), DAPT (10µM, Stem Cell Technologies), and Nicotinamide (NIC, 10 mM, Millipore-Sigma). DSM and IDE2 were included from day 6, NIC on days 1-10, FSK on days 1-30, and DAPT on days 18-30. Differentiation was induced in 5% CO2 at 37°C.

### GCaMP6s^+^ hPSC line generation

GCaMP6s was cloned into a human AAVS1 homologous recombination plasmid containing a puromycin resistance gene (Addgene #88698) downstream of a CAG2 promoter using a commercial vendor (Twist Bioscience, San Francisco, CA) and the sequence was verified by next generation sequencing. A plasmid encoding Cas9 and an AAVS1 targeting gRNA were kindly donated by the Xitiz Chamling Laboratory (JHU). Plasmids were transformed in competent in One Shot Stbl3 Escherichia coli (Invitrogen, C7373-03) and purified using ZymoPURE plasmid Maxi-prep following manufacturers protocols, followed again by next generation sequencing confirmation. For transfection, PSCs were grown in Matrigel coated 24-well plates to 60%. On the day of transfection, reactions were prepared using 25 ul OptiMEM media (Thermo Fisher Scientific, Waltham, MA), 350ng Cas9/gRNA plasmid and 750ng GCaMP plasmid, and lipofectamine stem cell reagent (Thermo Fisher Scientific). The complex was added to designated wells and incubated at 37°C. After 48h, media was exchanged for mTESR1 plus containing puromycin (0.5ug/mL) until clonal selection. After several days, adherent cells were resuspended replated at single-cell density in Matrigel-coated 96 well plates with 5uM of belebbistatin containing 200µL of mTeSR1 media. Colonies were grown for 7-10d and then passaged into larger plates. We generated 15 independent colonies and evaluated 3 of them.

### RT-PCR

Total RNA isolation was extracted from single cell-derived clones using TRIzol™ Reagent (Life Technologies, Inc., Rockville, MD), with RNA purified by a chloroform-isopropanol method. RNA quantity and quality were measured using a NanoDrop 2000 Spectrophotometer (Thermo Fisher Scientific). Complementary DNA (cDNA) was synthesized from 1.0 µg total RNA by reverse transcription using an iScript cDNA synthesis kit (Bio-Rad, Hercules, CA). The synthesized cDNA was used as a template for PCR amplification using specific primers designed to amplify the following specific sense and antisense PCR primer used were as follows: GFP 5’-CGTCTATATCAAGGCCGACAAG-3’ and 5’-CTTCTTCCGTGTCCCTGTATTT-3’; β-actin sense primer 5’-ACCATG GATGATGATATCGC-3’ and antisense primer 5’-ACA TGG CTG GGG TGT TGA AG-3 (loading control). PCR conditions were 95°C for 3 min (denaturation) and 35 cycles of 95°C for 5 seconds, 58°C for 5 seconds, 72°C for 15 seconds followed by 72°C for 10 min. PCR products were resolved by 2% agarose gel electrophoresis, visualized by ethidium bromide staining and imaged using the Gel Doc image system (Bio-Rad).

### Western blot

Whole cell lysates were prepared by scraping cells in 100µL of ice-cold RIPA buffer containing 1 mM of PMSF, 1mM of sodium ortho vanadate, 1mM of NaF and 1% of protease inhibitor cocktail. The lysate was rotated for 1 hour at 4°C followed by centrifugation at 12000xg for 10min at 4°C to clear cellular debris. Protein concentrations were quantified using a Bradford protein assay kit (Bio-Rad). Total proteins (50μg) were mixed with 2x Laemmli loading sample buffer (Bio-Rad) with 1% β-mercaptoethanol (Bio-Rad) and the samples boiled for 10 minutes. The proteins were resolved on Mini-protein 12% TGX gels (Bio-Rad) and transferred into nitro-cellulose membranes (Bio-Rad). After the membranes were blocked in 1x Tris-buffered saline containing 0.05% Tween-20 (TBST) and 5% fat-free milk, the membrane was incubated with primary antibodies (Supplemental Table 1) slowly on a shaker overnight at 4°C in 1xTBST with 5% bovine serum albumin. The next day, the membranes were washed three times for 5mins with 1xTBST and were further incubated with the corresponding horseradish peroxidase-conjugated secondary antibody at 37°C for 1 hour and then washed three times with 1x TBST for 5 mins. Immunodetection was performed by incubating membrane with 1:1 ratio of enhanced chemiluminescence detecting reagents (Western Bright ECL, Advansta, San Jose, CA) according to the manufacturer’s instructions. The iBright CL1500 Imaging system (Thermo Fisher Scientific) was used to capture the chemiluminescence signals of the bands.

### GCaMP6s Adeno-Associated Virus

GCaMP6s was cloned into an AAV packaging plasmid (VectorBiolabs #1369, Malvern, PA) containing a CAG2 promoter using a commercial vendor (Twist Bioscience, San Francisco, CA) and the sequence was verified by next generation sequencing. Plasmids were transformed in competent in One Shot Stbl3 Escherichia coli (Invitrogen, C7373-03) and purified using ZymoPURE plasmid Maxi-prep following manufacturers protocols, followed again by next generation sequencing confirmation. AAV was produced by the Powell Gene Therapy Center Vector Core Lab (University of Florida). The genetic payload was packaged into the AAV^DGEDF^ capsid, an AAV-2 variant that was developed by Dr. Shannon Boye (University of Florida) through *in vivo* directed evolution in nonhuman primates. Virus was delivered in BSS at a titer of 2.5×10^13^vg/mL.

### Mouse ocular hypertensive glaucoma

Mice were anesthetized with intraperitoneal ketamine (50 mg/kg), xylazine (10 mg/kg), and acepromazine (2 mg/kg) and topical 0.5% proparacaine hydrochloride eye drops (Akorn Inc. Buffalo Grove, IL, USA). One eye underwent intracameral injection of a 50:50 mixture of 6μm and 1μm diameter microbeads (2μL of each bead size, Polybead Microspheres®, Polysciences, Inc., Warrington, PA), followed by 1μL of viscoelastic compound (10 mg/mL sodium hyaluronate, Healon, Advanced Medical Optics Inc., Santa Ana, CA). IOP was measured using a Tonolab (Icare Finland Oy, Vantaa, Finland) at days 1, 3, 7, 10, 14, 21, and 28 to ensure a >5mmHg pressure elevation occurred.

### Transmission electron microscopy

Animals for TEM were euthanized by exsanguination under general intraperitoneal anesthesia, followed by intracardiac perfusion for 3 minutes with 4% paraformaldehyde in 0.1 M sodium phosphate buffer (Na_3_PO_4_, pH = 7.2), 1 minute with 0.1M cacodylate buffer, and 7 minutes with 2% paraformaldehyde / 2.5% glutaraldehyde in cacodylate buffer. Eyes were enucleated and kept in fixative overnight. Tissue was post-fixed in 1% osmium tetroxide, dehydrated in ascending alcohol concentrations, and then stained with 1% uranyl acetate in 100% ethanol for 1 hour. Tissues were embedded parallel to the optic axis (longitudinal sections) in an epoxy resin mixture at 60°C for 48 hours. One micrometer thick sections were cut and stained with 1% toluidine blue. Ultra-thin sections (∼68nm thick) were collected on copper grids and stained with uranyl acetate and lead citrate before examination with a Hitachi H7600 transmission electron microscope (Hitachi High Technologies, Clarksburg, MD).

### Scanning electron microscopy

Retinal samples were fixed in 2.5% glutaraldehyde, with 3mM MgCl_2_, in 0.1 M sodium cacodylate buffer, pH 7.2 overnight at 4°C. After buffer rinse, samples were postfixed in 1% osmium tetroxide in 0.1 M sodium cacodylate buffer for 1h on ice in the dark. Following a DH_2_O rinse, samples were dehydrated in a graded series of ethanol and left to dry overnight (in a dessicator) with hexamethyldisilazane (HMDS). Samples were mounted on carbon coated stubs, coated with 15nm of gold-palladium (Denton Desk3) and imaged on a Helios focused ion beam SEM (Thermo Fisher Scientific).

### Tissue Preparation and Fixation for Confocal Microscopy

Animals were euthanized and eyes were enucleated and placed in a Petri dish containing D-PBS (Thermo Fisher Scientific). The anterior segment tissues were removed, and the retina was gently detached from the posterior eyecup. Four radial incisions were made in the retina, and it was flatmounted on 13mm MF-Millipore hydrophilic membrane filters (Millipore-Sigma) in Maltese cross configuration. The retina flatmount was then transferred into a 4% PFA (Electron Microscopy Sciences, Hatfield, USA) solution for one hour at 4°C, and then transferred into PBS at 4°C.

### Cryosectioning

Whole eyes were harvested and immerse-fixed in 4% PFA (Electron Microscopy Sciences, Hatfield, USA) at 4°C overnight. Globes were cryoprotected by serial immersion in 5% and 20% sucrose solutions (Millipore-Sigma) for 3 hours each. Globes were then embedded in optimal cutting temperature (OCT) (Sakura Finetek, Torrance, USA) compound and frozen on dry ice. Tissue blocks were stored at -80**°**C, until use. A cryostat (Leica Microsystems, Wetzlar, Germany) was used to cut transverse sections 18 μm thick, which were thaw-mounted on glass slides.

### Immunofluorescence

Retinal flatmounts were washed with PBS for 5 min and then simultaneously blocked and permeabilized in 10% normal goat serum (NGS, Millipore-Sigma) and 0.3% Triton X (Millipore-Sigma) in PBS for 1h at room temperature (RT). Retinal flatmounts were then incubated in primary antibody for 4 days at 4°C on a shaker. Secondary antibody incubations were conducted overnight at 4°C on a shaker. Nuclei were counterstained with DAPI (1:10,000; Millipore-Sigma). Tissue was cover slipped under Aqua-Poly Mount (Polysciences, Warrington, USA) and stored at 4**°**C in the dark.

Retinal cryosections were washed with PBS and then then blocked and permeabilized in 10% normal goat serum (NGS, Millipore-Sigma) and 0.3% Triton X (Millipore-Sigma) in PBS for 1h at RT, prior to incubation in primary antibody for 24h at 4°C (Supplemental Table 1). The sections were then washed with PBS and incubated with corresponding secondary antibody for 1h at RT. Nuclei were counterstained with DAPI (1:10,000; Millipore-Sigma). Slides were cover slipped using Aqua-Poly Mount (Polysciences).

**Supplemental Table 1.**
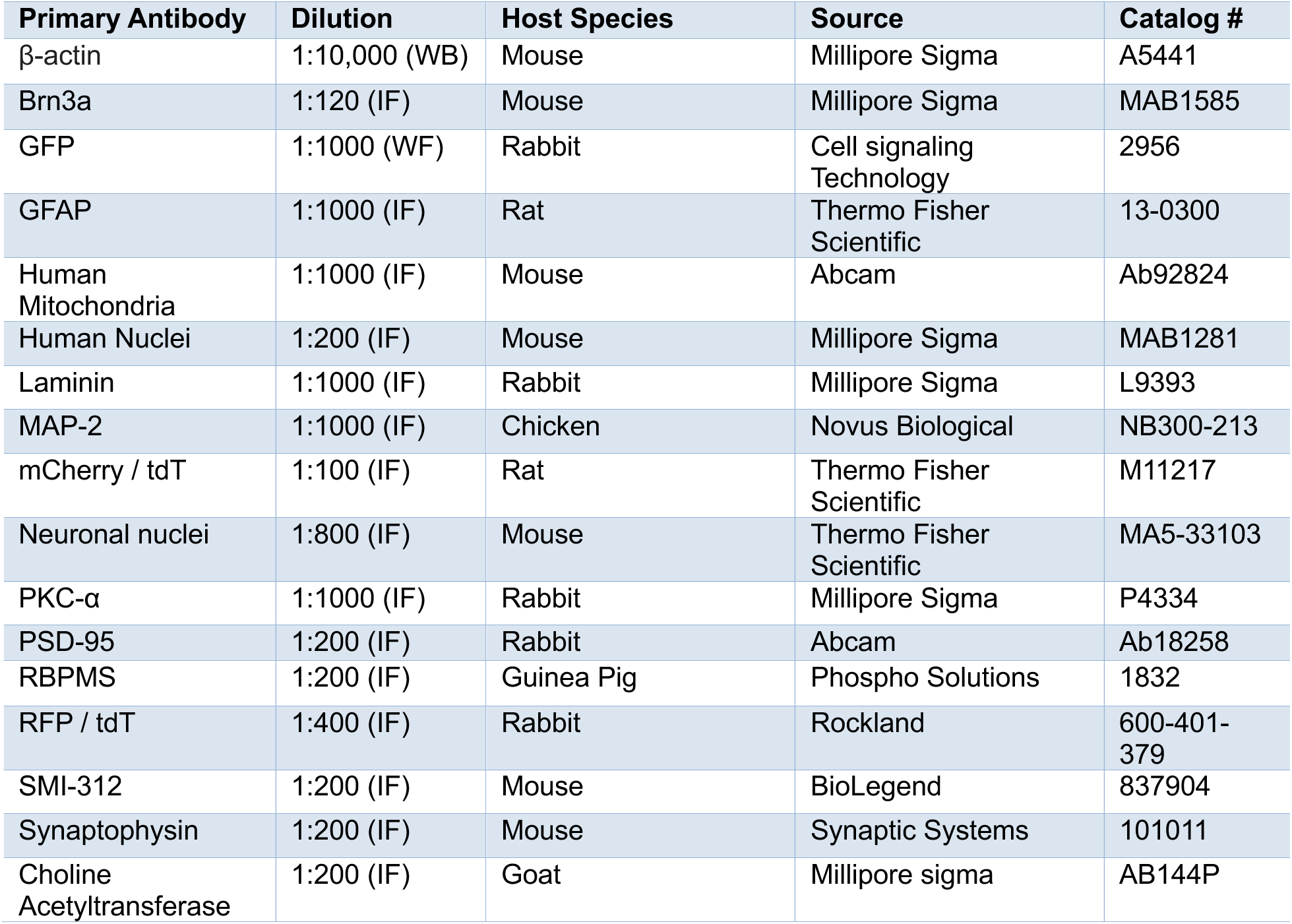
Primary antibody details for immunofluorescence. WB, Western blot. IF, Immunofluorescence.

## SUPPLEMENTAL FIGURES

**Supplemental Figure 1.**
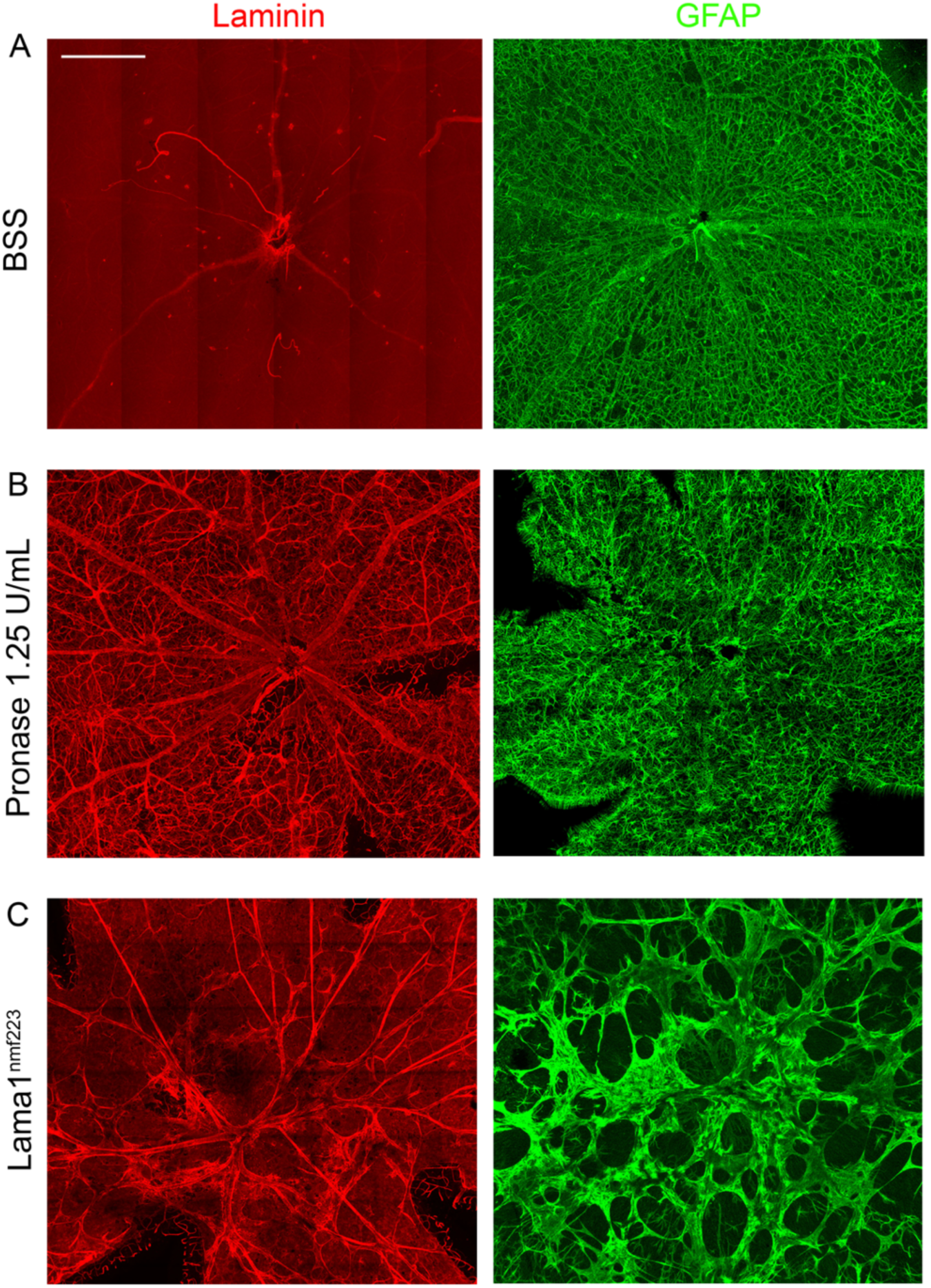
Effects of Pronase-E and the Lama1nmf223 mutation on retinal internal limiting membrane (ILM) and glia. (A) C57BL/6J mice were treated with intravitreal BSS and retinas were flatmounted 2 weeks later. (B) C57BL/6J mice were treated with intravitreal Pronase-E (1.25UmU and retinas were flatmounted 2 weeks later. (C) Lama1^nmf223^ mouse retinas were flatmounted and prepared for immunofluorescence. Tiled confocal z-stacks are shown, demonstrating immunofluorescence for laminin to highlight the ILM and vasculature (red) and GFAP to highlight astrocytes and Müller glia (green). BSS-treated animals had intact ILM and quiescent appearing glia. Pronase E was associated with ILM defects, which artifactually resulted in brighter labeling of the underlying vasculature due to enhanced antibody penetration. Pronase-E also was associated with glial hypertrophy. Lama1^nmf223^ retinas similarly demonstrated ILM defects but also exhibited preretinal gliovascular membranes. Scalebar, 1mm.

**Supplemental Figure 2.**
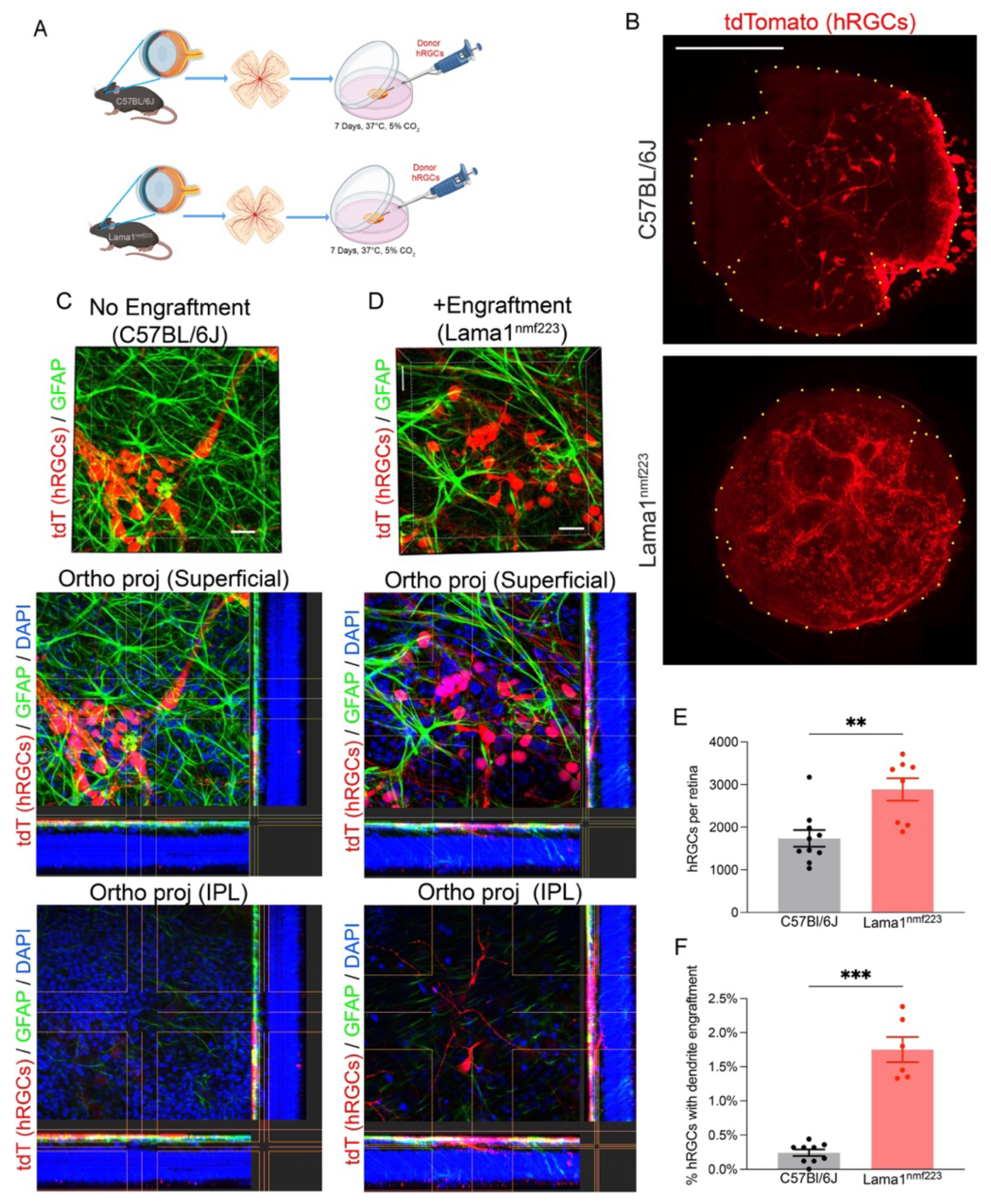
**Structural hRGC engraftment into wildtype and Lama1^nmf223^ organotypic retinal explant cultures (**A) Experimental design assessing hRGCs at 1 week following transplantation onto cultured organotypic C57BL/6J (with intact ILM) or Lama1^nmf223^ mouse retinal explants. (B) Example retinal flatmounts showing hRGC localization 1 week after co-culture. Scalebar, 1mm. (C) Example of donor hRGCs with somata and neurites superficial to the RGCL and GFAP^+^ astrocytes of the RNFL, with no processes visible within the IPL. Scalebar, 30 µm. (D) Example of donor hRGCs with somata co-planar to the host RGC layer (deep to GFAP^+^ astrocytes in the RNFL) and neurites extending into the IPL. Subcellular localization was evaluated using a multimodal assessment (see methods) Scalebar, 30 µm. Quantification of hRGC survival (E) and the rate of dendrite extension into the retinal parenchyma (F) within co-cultures.

**Supplemental Figure 3.**
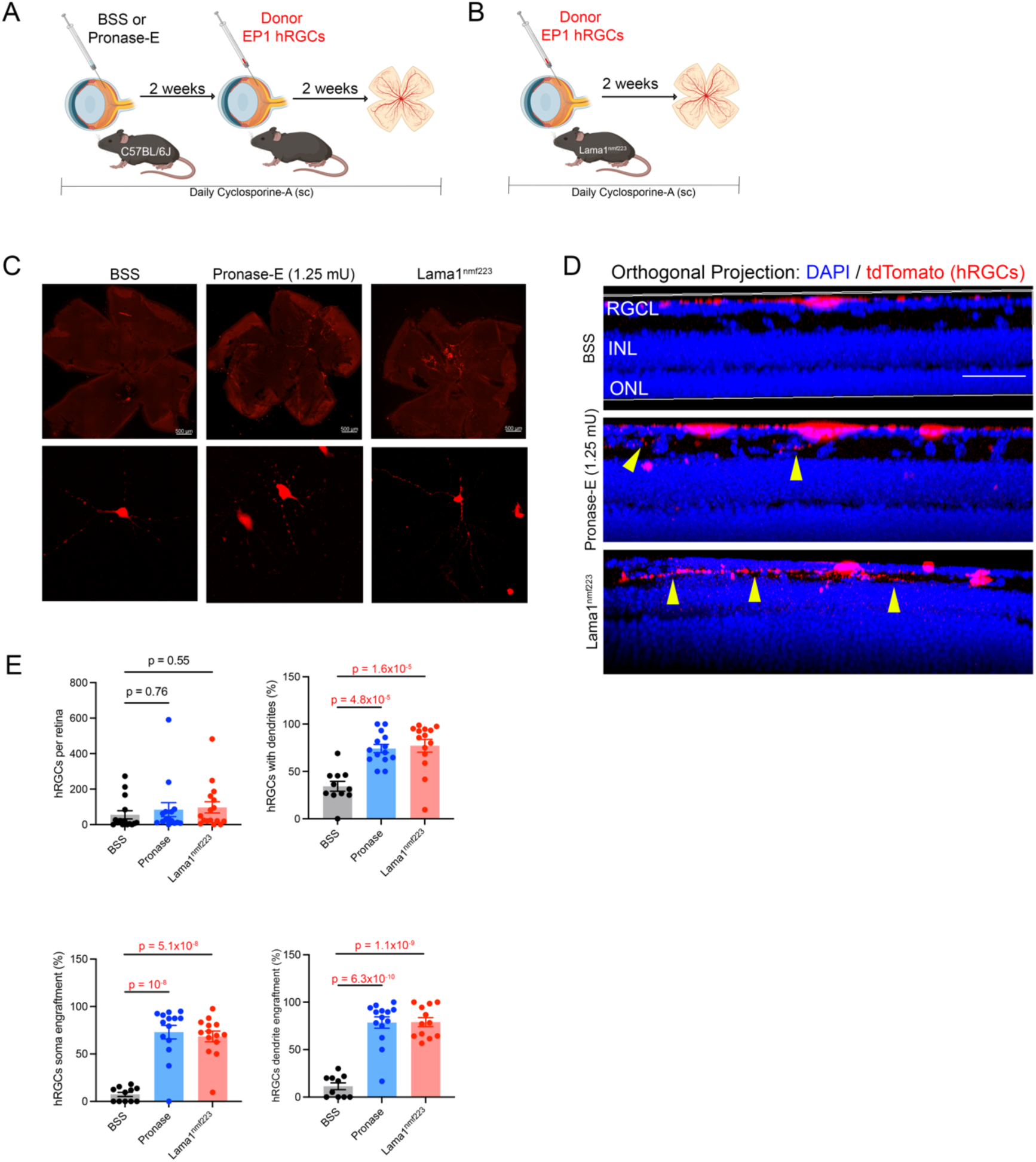
Structural EP1-hRGC engraftment in mice. hRGCs derived from EP1 human iPSCs were intravitreally injected in wildtype mice following pronase-E treatment (A) or directly into Lama1^nmf223^ mice (B). (C) Top row: retinal flatmounts demonstrating hRGC survival and localization 2 weeks after transplantation. Scalebar, 1mm. Bottom Row: high magnification images of donor RGCs extending neurites. (D) Orthogonal projections of confocal z-stacks demonstrating structural engraftment or engraftment failure. Arrowheads, hRGC dendrites within the inner plexiform layer (IPL). Scalebar, 20µm. (E) Quantification of hRGC survival, neurite extension, somal engraftment, and dendritic extension into the IPL.

**Supplemental Figure 4.**
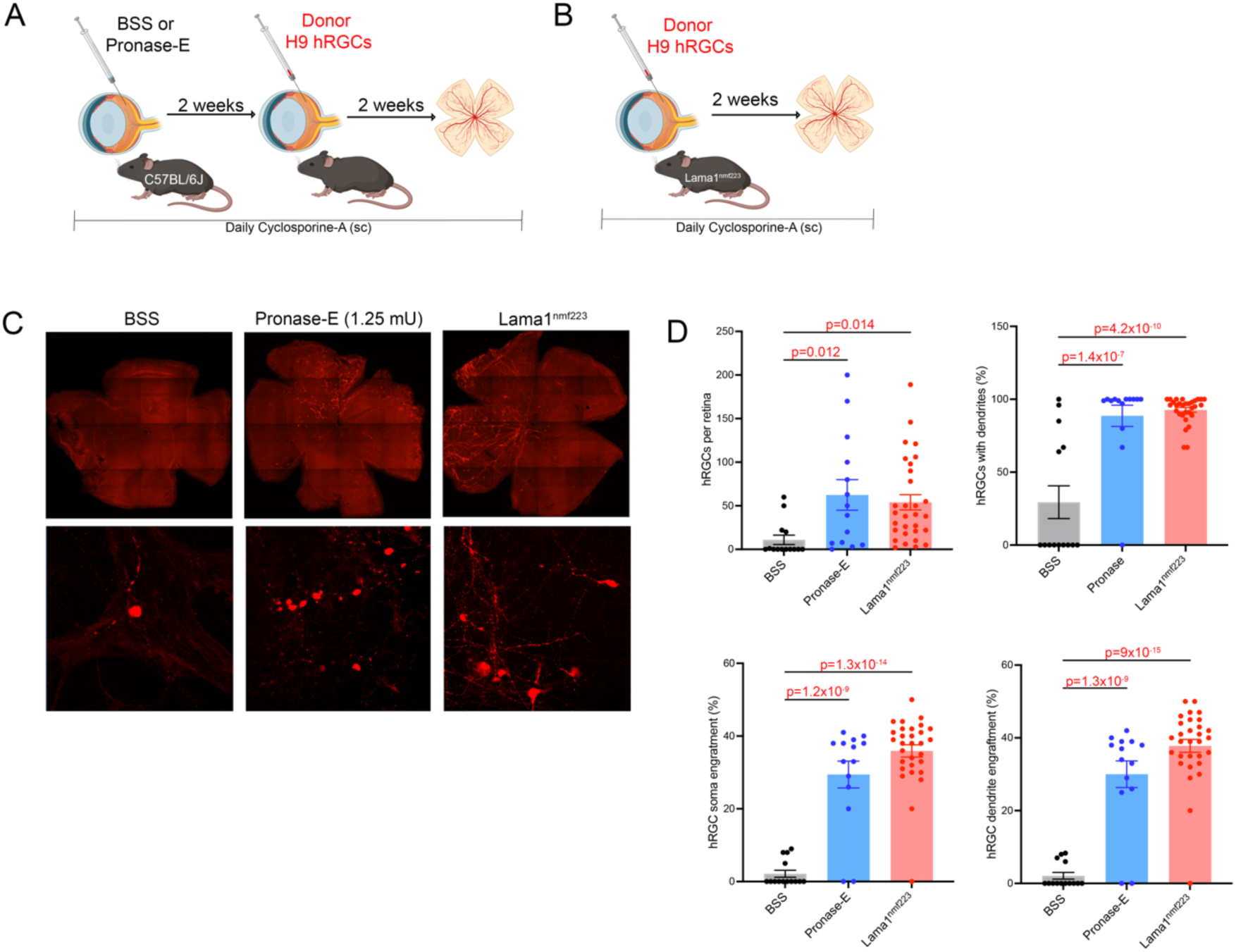
Structural H9-hRGC engraftment in mice. hRGCs derived from H9 human ESCs were intravitreally injected in wildtype mice following pronase-E treatment (A) or directly into Lama1^nmf223^ mice (B). (C) Top row: retinal flatmounts demonstrating hRGC survival and localization 2 weeks after transplantation. Scalebar, 1mm. Bottom Row: high magnification images of donor RGCs extending neurites. (D) Quantification of hRGC survival, neurite extension, somal engraftment, and dendritic extension into the IPL.

**Supplemental Figure 5.**
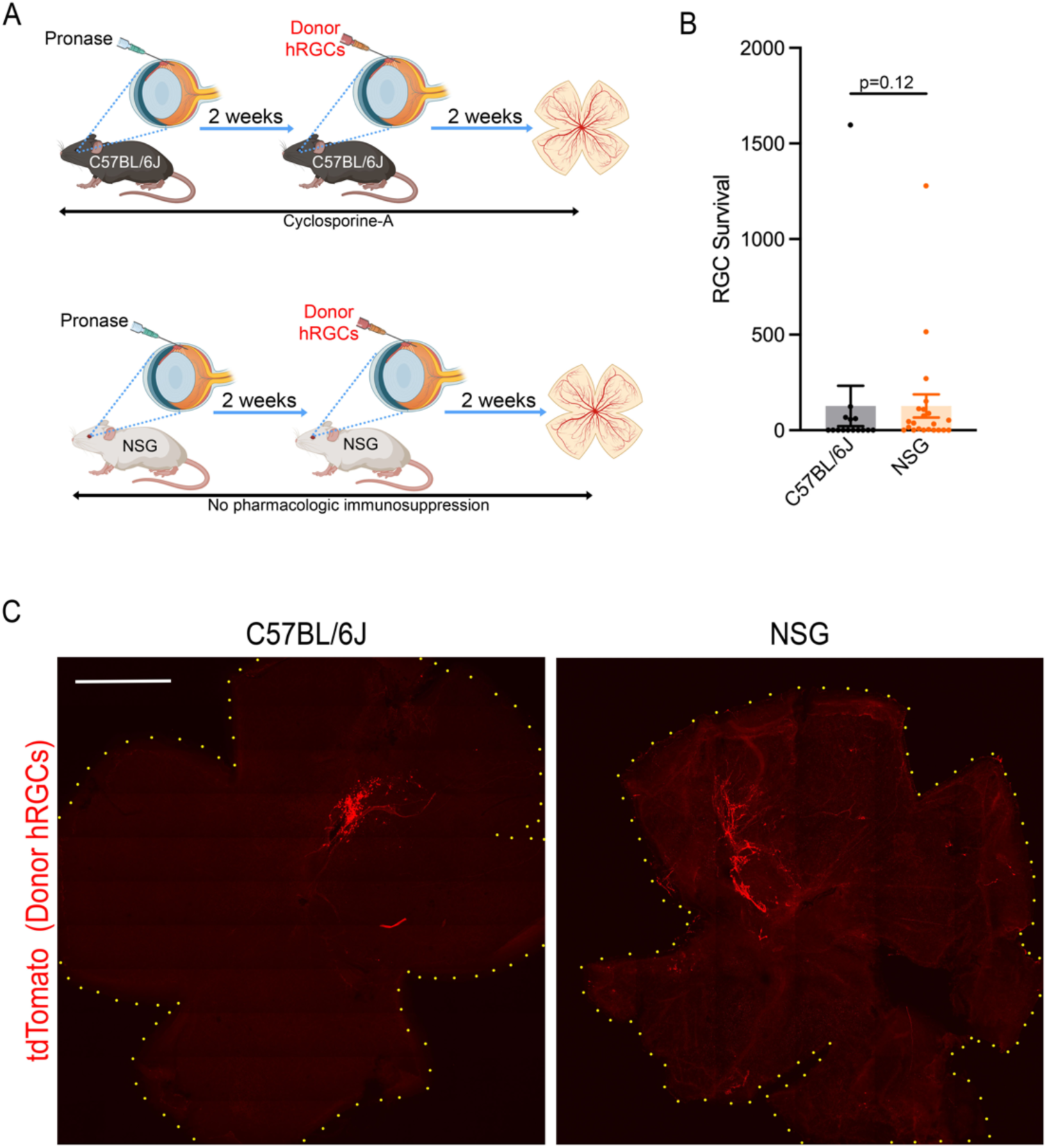
Survival of H7-hRGCs in immunosuppressed and immunocompromised mice. (A) hRGCs derived from H7 human ESCs were intravitreally injected in immunosuppressed C57BL/6J mice for NSG mice following pronase-E treatment. (B) Quantification of hRGC survival at 2 weeks following transplantation. (C) Retinal flatmounts demonstrating hRGC survival and localization 2 weeks after transplantation. Scalebar, 1mm.

**Supplemental Figure 6.**
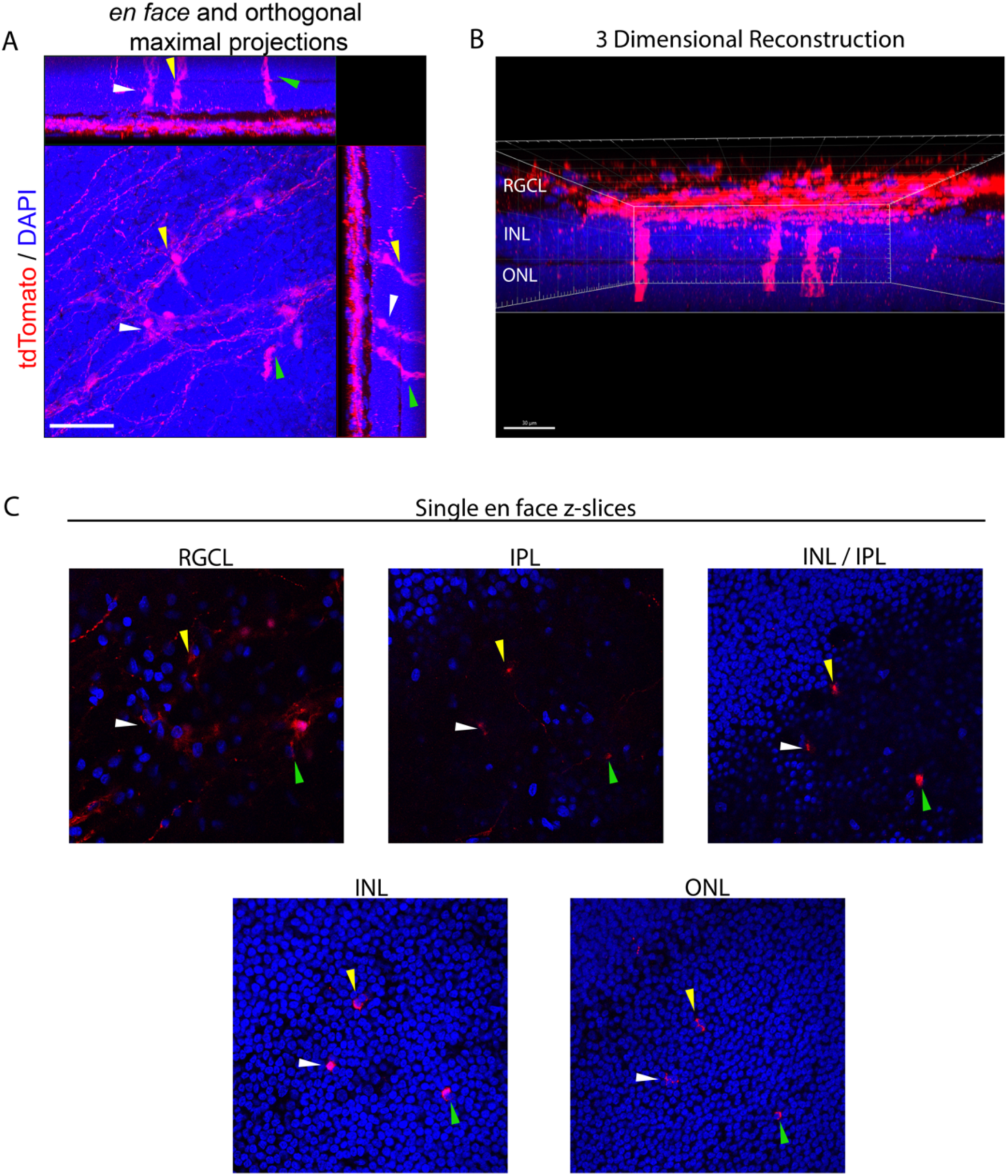
tdTomato intercellular material transfer to Müller glia following transplantation in nonhuman primates. (A) At 8 weeks following transplantation, confocal z-stacks and orthogonal maximum projections demonstrated tdTomato expression in donor RGCs and also in endogenous Müller glia (arrowheads). (B) 3D reconstruction of z-stack shown in panel A. (C) *En face* confocal slices taken through the retinal ganglion cell layer (RGCL), inner plexiform layer (IPL), inner nuclear layer (INL), and outer nuclear layer (ONL). Colors of arrowheads correspond to individual tdTomato^+^ Müller glia across images and panels. Scalebars, 30µm.

## SUPPLEMENTAL VIDEOS

**Supplemental Video 1. Volumetric demonstration of hRGC engraftment in a Pronase-E treated retina.** Two weeks following hRGC transplantation, retina was isolated and processed for confocal microscopy. A high-resolution z-stack was rendered in 3D. hRGCs express tdTomato (red) and nuclei are labeled with DAPI (blue) enabling visualization of the retinal layers. An hRGC which demonstrates somal localization to the endogenous RGCL and dendrite lamination within the IPL is traced (grey) to enhance visualization. Frames from 0:06 to 0:11 were cropped to remove the most superficial slices and enhance visualization of the deeper lying hRGC.

**Supplemental Video 2. Volumetric demonstration of hRGC engraftment in a Lama1nmf223 retina.** Two weeks following hRGC transplantation, retina was isolated and processed for confocal microscopy. A high-resolution z-stack was rendered in 3D. hRGCs express tdTomato (red) and nuclei are labeled with DAPI (blue) enabling visualization of the retinal layers. An hRGC which demonstrates somal localization to the endogenous RGCL and dendrite lamination within the IPL is traced (grey) to enhance visualization. Frames from 0:09 to 0:18 were cropped to remove the most superficial slices and enhance visualization of the deeper lying hRGC.

**Supplemental Video 3. Volumetric demonstration of hRGC engraftment in a nonhuman primate.** Eight weeks following hRGC transplantation, retina was isolated and processed for confocal microscopy. A high-resolution z-stack was rendered in 3D. hRGCs express tdTomato (red) and nuclei are labeled with DAPI (blue) enabling visualization of the retinal layers. hRGCs demonstrate somal localization to the endogenous RGCL and extensive dendrite lamination within the IPL, and occasionally deeper.

**Supplemental Video 4. tdTomato intercellular material transfer to Müller glia following transplantation in a nonhuman primate.** Eight weeks following hRGC transplantation, retina was isolated and processed for confocal microscopy. A high-resolution z-stack was rendered in 3D. hRGCs express tdTomato (red) and nuclei are labeled with DAPI (blue) enabling visualization of the retinal layers. hRGCs demonstrate somal localization to the endogenous RGCL and extensive dendrite lamination within the IPL, and occasionally deeper. In addition, three endogenous Müller glia are brightly tdTomato^+^.

**Supplemental Video 5. Volumetric evaluation of hRGC axonal extension into the optic nerve head.** Two weeks following hRGC transplantation, retina was isolated and processed for confocal microscopy. A high-resolution z-stack was centered on the optic nerve head and rendered in 3D. Axons were traced in grey to enhance visualization and noted to enter the optic nerve head.

**Supplemental Video 6. Calcium imaging of endogenous mouse RGCs following AAV transduction with GCaMP6s.** Two photon microscopy was used to capture a time-lapse series of images in a flatmounted retina. Spontaneous electrophysiological activity is noted throughout the time course (fluorescent flickering). A visible light stimulus was applied at 10sec, resulting in a transient increase in fluorescence in many, but not all, RGCs. Framerate, 14fps.

**Supplemental Video 7. Calcium imaging of endogenous mouse RGCs following AAV transduction with GCaMP6s.** Two photon microscopy was used to capture a time-lapse series of images in a flatmounted retina. A train of visible light stimuli were applied every 10 seconds, resulting in a transient increase in fluorescence in many, but not all, RGCs that decreased in intensity over multiple stimulations. Framerate, 7fps.

**Supplemental Video 8. Calcium imaging of endogenous mouse RGCs following AAV transduction with GCaMP6s, with heatmap visualization.** Supplementary video 7 was rendered using a fluorescent intensity heatmap to better visualize changes in fluorescent intensity. Refer to Fig 8I for scale.

**Supplemental Video 9. Calcium imaging of transplanted hRGCs in Lama1^nmf223^ retina.** Two photon microscopy was used to capture a time-lapse series of images in a flatmounted retina. Spontaneous electrophysiological activity is noted throughout the time course (fluorescent flickering). A train of visible light stimuli were applied every 10sec (indicated by “STIMULATION” overlay on the top right). A subset of hRGCs responded to simulation with a transient increase in GCaMP6s fluorescence, though many hRGCs did not respond to each stimulation in the series. Framerate, 7fps.

**Supplemental Video 10. Calcium imaging of transplanted hRGCs in Lama1^nmf223^ retina, with heatmap visualization.** Supplementary video 9 was rendered using a fluorescent intensity heatmap to better visualize changes in fluorescent intensity. Refer to Fig 8I for scale.

**Supplemental Video 11. Calcium imaging of a transplanted hRGC in Lama1^nmf223^ retina.** Two photon microscopy was used to capture a time-lapse series of images in a flatmounted retina. Spontaneous electrophysiological activity less prominent in this cell than in those from Supplemental Video 9-10. A train of visible light stimuli were applied every 10sec. A single visualized hRGC responded to simulation with a transient increase in GCaMP6s fluorescence. Framerate, 7fps.

**Supplemental Video 12. Calcium imaging of transplanted hRGCs in Lama1^nmf223^ retina, with heatmap visualization.** Supplementary video 11 was rendered using a fluorescent intensity heatmap to better visualize changes in fluorescent intensity. Refer to Fig 8I for scale.

## Notes

### Competing Interest Statement

The authors have declared no competing interest.

### Summary of Updates

We have added data from additional experiments including RGC transplantation in rats, a mouse model of glaucoma, and non-human primates. We have also added data from calcium imaging.

